# Designer DX-tile DNAns hydrogels

**DOI:** 10.1101/2025.03.05.641269

**Authors:** Dylan V. Scarton, Alessandra B. Coogan, Eray O. Tulun, Katie A. Harrison, Peter M. Touma, Jack Buchen, Richard C. Steiner, Christopher R. Fellin, Hunter G. Mason, Chih-Hsiang Hu, Xiaoning Yuan, Shailly Jariwala, Remi Veneziano

## Abstract

Pure deoxyribonucleic acid (DNA) hydrogels synthesized via the hybridization of multi-arm DNA tiles are uniquely programmable and functionalizable biomaterials suitable for applications ranging from biosensing to cell-free protein production and soft tissue engineering. However, the full potential of the design flexibility and functionalization offered by DNA molecules has not yet been leveraged for pure DNA hydrogels, thereby limiting their range of mechanical properties and reducing their versatility and broader use. In this study, we introduce multi-arm double-crossover (DX)-tile motifs, often used in DNA nanoparticle design, to enable greater control over the hydrogel’s mechanical properties and facilitate functionalization. Specifically, we demonstrate that modifying structural design parameters, such as the arm geometry, length, valency, and linker design, allows fine control of the elastic modulus and viscoelastic properties of the hydrogels. We also show that functionalization can be performed without compromising the hydrogels’ physical properties and exhibit enhanced mechanical strength and tunable properties, compared to simple duplex-based DNA hydrogels. Furthermore, these DNA hydrogels demonstrated printability and scalability, which pave the way towards the development of novel formulations and bioinks for the rational design of soft tissue engineering scaffolds and broaden the use of DNA hydrogels for other biomedical applications.

## Main

Deoxyribonucleic acid (DNA) molecules have unique structural, mechanical, and biochemical attributes that can be leveraged to assemble biomaterials from the nanoscale to the macroscale with tunable physical properties and precise functionalization^1–3^. Notably, DNA molecules can be used to synthesize DNA-based hydrogels or pure DNA hydrogels. DNA-based hydrogels are generally composed of polymers such as polyethylene glycol diacrylate^4^, ethylene diglycidyl ether^5,6^, or oxidized alginate^7^ that are crosslinked with small DNA duplexes or motifs like G-quadruplexes^8^ and i-motifs^9^ to create dynamic three-dimensional (3D) polymeric meshes^10^. These hybrid hydrogels have been successfully used as responsive biomaterial scaffolds for 3D cell culture^11^, tissue engineering^12^, and soft robotics^13^ leveraging their mechanical properties that can be controlled via tuning the cross-linking ratio and the nature of the DNA linker used^14–16^. Pure DNA hydrogels, on the other hand, are exclusively comprised of nucleic acid molecules (e.g., 2D motifs or long rolling circle amplification strands^17,18^) that act as building blocks to form 3D polymeric meshes^19^. Among them, multi-arm tile hydrogels are one interesting type compared to the other DNA-based hydrogels^20,21^. These hydrogels, which are formed via the hybridization of two-dimensional (2D) single duplex-based DNA motifs bearing complementary overhangs^22,23^, offer superior programmability, addressability, and design flexibility.

The individual motifs, also known as DNA nanostars (DNAns), typically feature three arms that each comprise a single DNA duplex with arm lengths of 20 nucleotides (nts) for a diameter of roughly seven nanometers (nm)^24^. Their mechanical properties are tuned by controlling the concentration of motifs or adjusting the number of complementary base pairs in the linker. In some cases, exploiting the stimuli-responsive nature of DNA allows for reorganization of the crosslinking network at room temperature^25–27^. However, the design simplicity and limited crosslinking capabilities of these multi-arm hydrogels have restricted their range of mechanical properties,^23,28^ resulting in low elastic moduli ranging from only five to a few hundred pascals (Pa)^29,30^, due to the inherent flexibility of DNA molecules and fewer weak hydrogen-bonding crosslinks^31,32^. Thus, despite nearly two decades of advancements in DNA nanotechnology, pure DNA hydrogels have not yet been fully leveraged to their complete potential. They are largely restricted in their use to specific subsets of applications that do not require mechanical strength or complex functionalization, such as biosensing and drug delivery^33,34^.

In this study, we leveraged DNA nanotechnology design strategies commonly used for DNA nanoparticle synthesis to enhance the design flexibility of DNAns hydrogels and increase their range of mechanical properties and complex functionalization for subsequent applications like tissue engineering. Specifically, we replaced the single-duplex-based arm in the DNA motifs with double-crossover (DX)-tile motifs, common in constructing wireframe DNA origami nanostructures^35^, to create DX-DNAns hydrogels. DX-tile motifs offer several advantages, such as increased stiffness, more functionalization sites, and improved design flexibility^36^. Our results demonstrated that the DX-DNAns hydrogels have tunable elastic moduli from ∼0.1 kPa to more than 1.2 kPa and possess viscoelastic properties conducive to printability. Importantly, our design strategy enabled this tunability and ease of functionalization without compromising its structural integrity, thereby highlighting its potential to serve as a design platform for ‘smart’ biomaterials.

### Design and formation of DX-tile DNAns hydrogels

To explore the influence of DX-tile-based DNA motifs on the physical properties of DNA hydrogels and facilitate comparison with previously published DNAns hydrogels, we first developed multiple variants of a three-way junction (3WJ), comprised of a central core and three identical arms (***Figure 1a****, left*; ***Figure S1a***). Each 3WJ structure is assembled with multiple short oligonucleotide strands that are used in various ratios (***Table S1***). This synthesis strategy increased cost-effectiveness by reusing different oligonucleotide strands in bulk to explore various arm geometry combinations. To enable hybridization of multiple motifs, each 3WJ arm was modified with two short single-stranded overhang linkers (one linker per duplex) that paired with complementary sequences of the same length on another 3WJ motif (***Table 1***; ***Figure S2***). To examine the role of linker properties on downstream hydrogel behavior, we designed three different linker types: one short seven-base single strand overhang (Sh), one long 21-base single strand overhang (Lo), and one variant of the Lo linker with a non-hybridizing region of seven bases in the middle of its sequence called mismatch linker (Lm) (***Figure 1a****, right*). We also designed a blunt end (Bl) version of the 3WJ that did not bear any ssDNA linkers to assess the role of base stacking in gel formation, as typically observed for DNA nanoparticles^37,38^. For all the 3WJ constructs, our models showed arm-to-arm length without the linker to be estimated at 31.6 nm with arm angles of 118.2 degrees (***Figure S3***). After validating the sequences with NCBI BLAST to avoid mispairings, the different parts of the 3WJ and the entire structures were slowly annealed, and their folding was checked with agarose gel electrophoresis. The gel image (***Figure S4***) demonstrates a monodispersed 3WJ core motif and assembly of the three complete structures (with Sh, Lo, and Lm linkers) with limited byproducts. To characterize the assembly of the different core and arm parts of the structures, we used a fluorescence resonance energy transfer (FRET)-based experiment wherein a strand was modified with a Cy3 dye and another with a quencher. Moving the location of the Cy3 dye and quencher validated the incorporation of all strands and the stepwise formation of the 3WJ motifs, with the arm flexibility affecting the FRET signal (***Figure S5)***. With atomic force microscopy (AFM), we confirmed proper 3WJ folding (***Figure 1b***). Finally, with dynamic light scattering (DLS), we measured the hydrodynamic diameters of each individual motif showing an average of ∼51.0 ± 6.7 nm for 3WJ-Sh, ∼60.6 ± 1.0 nm for 3WJ-Lo, and ∼75.1 ± 11.0 nm for 3WJ-Lm, which all appear congruent with the estimated values (***Figure 1c***). Subsequently, we determined the melting temperatures of the motifs via quantitative polymerase chain reaction (qPCR) to check their stability at physiological temperatures (***Figure 1d***). The core and each arm have similar melting temperatures at 65°C and 64°C, respectively, which is typical for DX-tile-based nanoparticles^35^. For the full motifs, two peaks are visible at 64°C and 80°C due to the enhanced thermal stability induced by the cooperative assembly of the core and the arms, which confirmed the formation of the entire structure. Overall, our results demonstrated the proper formation of all 3WJ motifs designed.

**Figure 1.**
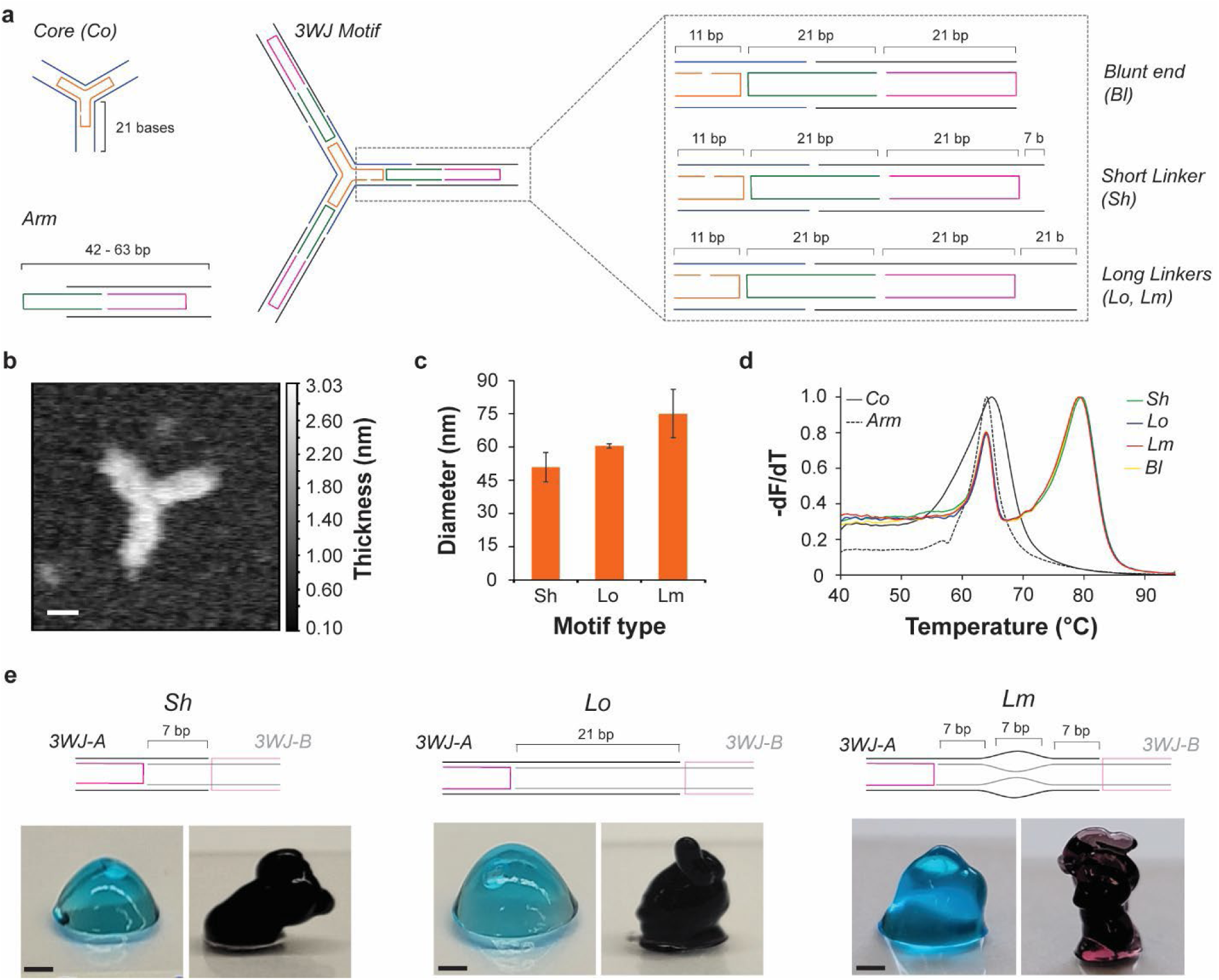
DX-tile-based DNA motif folding and DX-DNAns hydrogel formation for 3WJ constructs. (a) Initial design of the DX-DNAns constructs, showing the core and arm elements (*left*), and the different arm linkers (*right*). (b) Representative atomic force microscopy image showing proper folding of a 3WJ-Lo motif (scale bar: 10 nm). (c) Average diameter of 3WJ-motifs with different linkers, as measured by dynamic light scattering. Error bars represent standard deviation of the mean (n = 3 independent samples/group). (d) Melting curves of motif constituents obtained by qPCR (n = 4 independent samples/group). (e) Representative array images of hydrogels assembled with motifs bearing different linker types and at different motif concentrations (*60 µM, blue; 90 µM, black*; scale bars: 1 mm).

**Table 1.** Measured storage moduli (G’) for the hydrogels assembled with different 3WJ and 6WJ constructs. Standard deviation shown as ± (*n* = 3 independent samples/group).

| Name | 3WJ |  | 6WJ |  |
| --- | --- | --- | --- | --- |
| [motifs] | $60\text{ }\mu\text{M}$ (Pa) | $90\text{ }\mu\text{M}$ (Pa) | $25\text{ }\mu\text{M}$ (Pa) | $50\text{ }\mu\text{M}$ (Pa) |
| 3/6WJ-Sh | $84 \pm 11$ | $150 \pm 31$ | $119 \pm 12$ | $1266 \pm 230$ |
| 3/6WJ-Lo | $135 \pm 11$ | $410 \pm 130$ | $179 \pm 46$ | $1095 \pm 8$ |
| 3/6WJ-Lm | $101 \pm 13$ | $255 \pm 13$ | $282 \pm 25$ | $1201 \pm 128$ |

We then tested hydrogel formation by mixing complementary motifs at equimolar concentration ranging from 15 to 90 µM (***Figure 1e***, *top*). At 15 µM, only the solutions containing complementary motifs with Lm linker formed hydrogels, while other motifs remained liquid, flowing on a tilted glass surface (***Figure S6***). At 30 µM, all tested motifs formed and held shape (***Figures S6, S7***). At higher concentrations, hydrogels clearly formed for all motifs, as shown in ***Figure 1e*** for hydrogels formed with the Sh (*left*), Lo (*middle*), and Lm (*right*) linkers at concentrations of 60 µM (*blue*) and 90 µM (*black*). The 90 µM samples consistently appeared thicker than 60 µM across the three linker types tested. Notably, although these results could be attributed to an increased concentration of the DNA motifs, a solution of a single motif type with overhangs at 60 µM or non-complementary motifs mixed at 60 µM did not yield hydrogel formation (***Figure S8***). Single Bl motif mixtures, on the other hand, formed a hydrogel directly after folding of the individual motifs, as opposed to the other motif types that required mixing complementary motifs, likely due to blunt-end stacking (***Figures S9)***^39,40^.

We used scanning electron microscopy (SEM) to further characterize the hydrogel microstructure. The 30 µM hydrogels exhibited a very loose fiber-like mesh, likely due to the low concentration of motifs, reducing its crosslinking ability (***Figures S10a, S11***). In contrast, the 60 and 90 µM hydrogels, formed regular and uniform microporous mesh networks with smooth, round pores that are typically observed in other polymeric and pure DNA hydrogels^41–43^ (***Figure 2a***; ***Figures S10b,c***). To determine if the linker type influenced porosity, we systematically measured the pore sizes for each hydrogel (***Figure S12***), which ranged from ∼4 to 6 µm in diameter (***Figure 2b***; ***Figure S13***) across all three linker types. The concentration ranges did not significantly affect the pore size for all motifs, but small and significant variations were observed between the different motif types at 60 µM (3WJ-Sh: 4.4 ± 0.2 µm; 3WJ-Lo: 5.0 ± 0.3 µm; 3WJ- Lm: 5.5 ± 0.2 µm) and 90 µM (3WJ-Sh: 4.8 ± 0.2 µm; 3WJ-Lo: 5.4 ± 0.3 µm; 3WJ-Lm: 5.7 ± 0.2 µm). Of note, hydrogels formed with the 3WJ-Bl motifs displayed similar pore sizes (5.8 ± 0.3 µm and 5.3 ± 0.3 µm for 60 and 90 µM, respectively). These results suggest that the micro-pore sizes were only marginally affected by the linker types within the range of motif concentrations tested. Furthermore, AFM imaging of the hydrogels (***Figure 2c*** and ***Figure S14***) revealed the presence of round and regular nano-pores with smooth edges created from the nanoscale assembly of the motifs, as previously observed with DNA hydrogels^44,45^. . Taken together, these results demonstrate that DX-tile motifs can be used to form hydrogels. All tested linkers formed hydrogels at the minimum concentration of 30 µM, including the base-stacking crosslinks of the Bl arm terminus that formed with similar structural properties.

**Figure 2.**
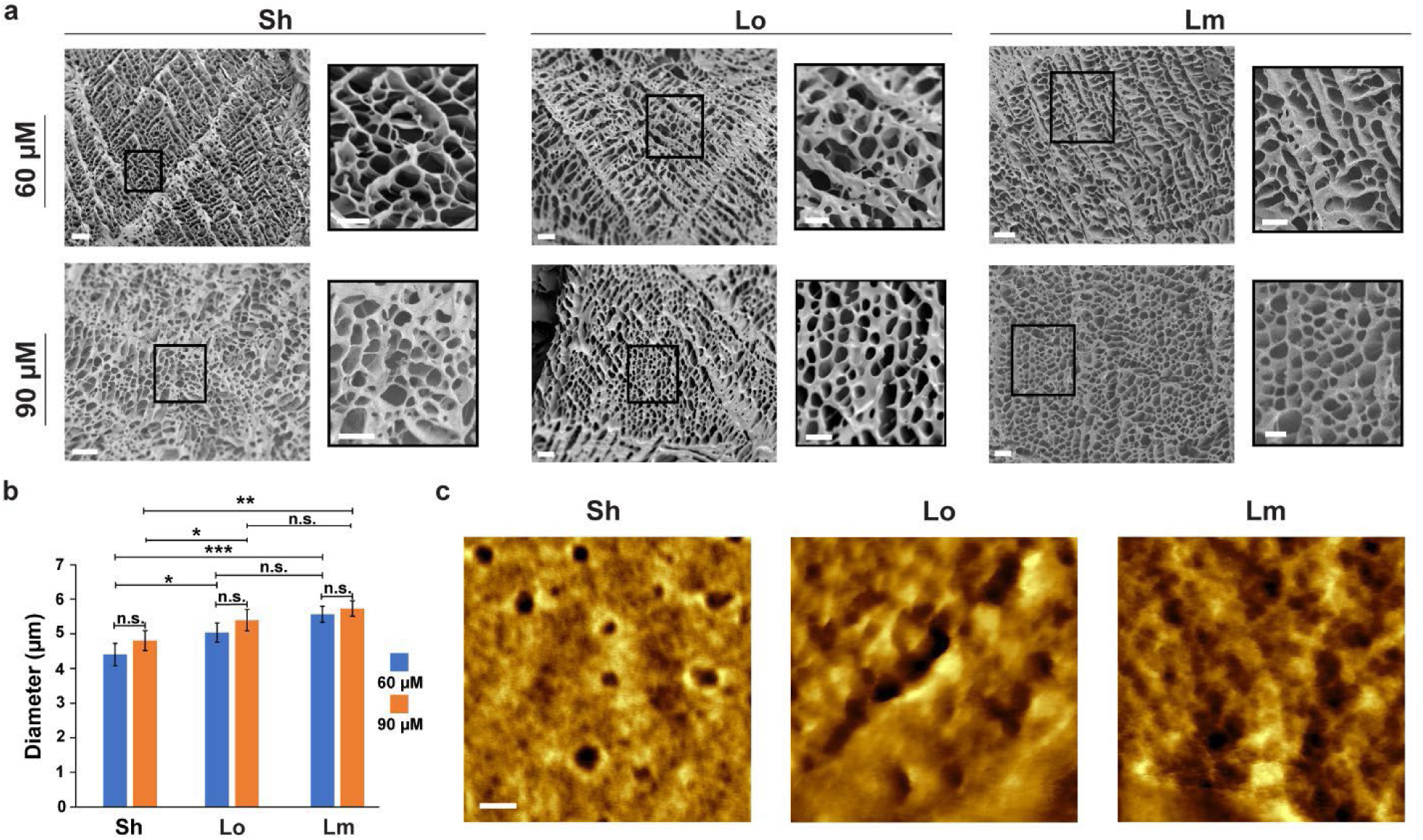
Structural characterization of the 3WJ hydrogels. (a) SEM images of the dried hydrogels (scale bar: 20 µm) with emphasis on the micro-pore morphology (scale bar: 10 µm) (b) Micro-pore size comparison across the various 3WJ hydrogels assembled with different linkers. Error bars represent standard error of the mean (*n* = 50 independent samples/group). *P*-values were calculated by student t-test (not significant, n.s.; \**p*<0.1, \*\**p*<0.01, \*\*\**p*<0.001). (c) Nano-pore morphology observed by AFM (scale bar: 50 nm).

### Nano-structural features and motif concentration alter bulk mechanical properties

Next, we investigated the effects of the design elements for the DX-tile nanostructure on the bulk viscoelasticity and mechanical strength of the hydrogels via rheological testing^46^. For comparison with single duplex motifs typically used for DNAns hydrogels, we designed three-armed single duplex motifs (Y motifs) with 7- (Y7) and 21- (Y21) base overhang linkers (***Table S2 and Figure S15***). The proper folding and monodispersity of the two Y motifs were confirmed by agarose gel electrophoresis (***Figure S16a***). Hydrogel formation was verified both visually (***Figure S16b***) and with SEM (***Figures S16c***) for motif concentrations ranging from 37.5 to 225 µM for Y7 and 30 to 180 µM for Y21 to approximate some of the motif concentrations used for the DX-DNAns hydrogels and the total DNA concentration (***Table S3***).

We first performed amplitude sweeps (***Figure 3a***) to establish the linear viscoelastic range (LVER) and define the extent to which the hydrogel can be deformed under controlled oscillatory strain before yielding. The storage modulus (G’) represents the elastic, solid-like behavior of the material, while the loss modulus (G”) represents its viscous, liquid-like behavior. The experimentally-determined modulus crossover or flow point is the strain at which the material transitions from dominant solid-like behavior (G’ > G”) to dominant liquid-like behavior (G” > G’). The DX-DNAns hydrogels assembled with the Sh linker exhibited the highest modulus crossover points (30% at 60 µM; 90% at 90 µM) and those with the Lo linker (3% at 60 µM; 20% at 90 µM) had the lowest crossover points, while those of the Lm linker hydrogels fell in-between (12% at 60 µM; 30% at 90 µM). This result indicates that the Sh linker allowed for more elasticity, likely due to the more transient nature of hybridization of the 7-bp linker over the 21-bp linker. Overall, increasing the motif concentration duly increased the crossover point of the hydrogels. This result is consistent with ranges observed for single-duplex DNA hydrogels between ∼10-80%^47^ and similar DNA hydrogels between ∼6-50%^15^, as determined by material concentration and therefore hydrogel stiffness^15^. No definitive LVER was observed for the Y-motif hydrogels at a matching motif concentration of 60 µM (***Figure S17a, top***), but, notably, when evaluated at motif concentrations approximately matching the DNA content to the 3WJ hydrogels, the concentration-dependent trend in the crossover point was inverted (***Figure S17b, top***). The flow points occurred at ∼83% and ∼104% strain (*p* = 0.084) for Y7-150 µM and Y21-120 µM, respectively, suggesting both nanostructure and linker design influence the LVER^15^. The LVER for these Y-motif-based hydrogels was shorter than those of the DX-tile hydrogels at a matching motif concentration and longer for the Y-motif-based hydrogels made with the same DNA concentration, which is expected because the DX-tile motifs require more energy to dehybridize their two linkers. As a control, we performed amplitude sweeps on solutions of the single 3WJ-Sh motif at 60 µM (***Figure S18a***) that did not yield hydrogels (***Figure S8***). As expected with non-hybridized DNA motifs, we did not observe a clear LVER, as the moduli inverted at multiple points and the plot showed considerable variance consistent with a non-gel material. The rheological profiles of the DX-DNAns hydrogels formed at 30 µM (***Figure S19a***), while more viscous than the singular motif, possessed non-definitive LVERs and higher crossover points (118% for 3WJ-Sh; 49% for 3WJ-Lo; 109% for 3WJ-Lm) than those of the 60 and 90 µM hydrogels (***Figure 3a***). This behavior is likely due to its stretched fiber-like organization and defects also observed by SEM (***Figures S10a and 11***).

**Figure 3.**
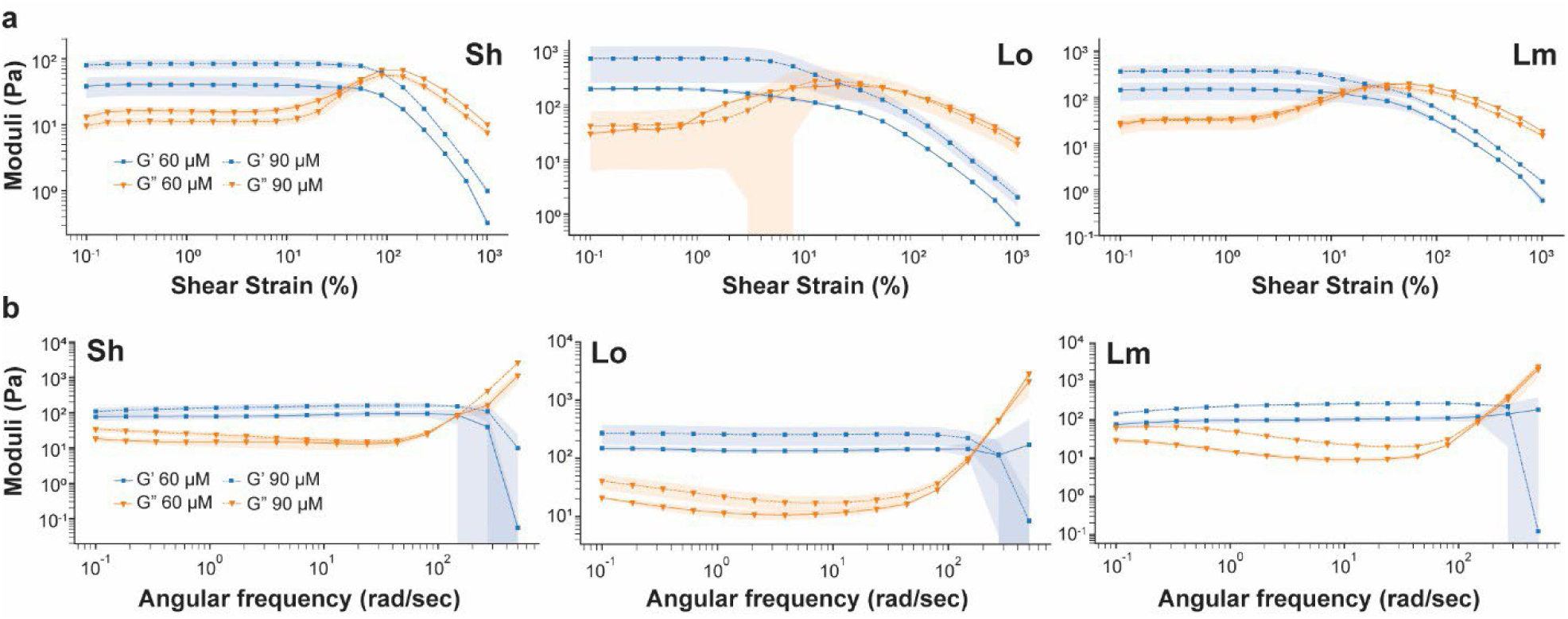
Rheological characterization profiles of the 3WJ constructs. (a) Amplitude sweeps of the hydrogels assembled at 60 µM and 90 µM of motifs with three different linkers. Error shadings represent standard deviation of the mean (*n* = 3 independent samples/group). (b) Frequency sweeps of the hydrogels assembled at 60 µM and 90 µM of motifs with three different linkers. Error shadings represent standard deviation of the mean (*n* = 3 independent samples/group).

Following those characterizations, we conducted frequency sweeps (***Figure 3b***) to capture the frequency-dependent rheological properties of the hydrogels. We observed distinct rheological profiles emerging across all the formulations and conditions tested, compared to a standard of 1 hertz (Hz) (***Table 1***). For the 60 µM hydrogels, the Lo linker hydrogels had the highest G’ at 135 ± 11 Pa, while those formed with Sh linker had the lowest G’ at 84 ± 11 Pa and those of the Lm linker were in-between at 101 ± 13 Pa. This trend is the exact opposite of the amplitude sweep results, which suggests an inverse relationship of the LVER and the G’, as described in other work related to DNA hydrogels where greater rigidity of the DNA scaffold increased DNA hydrogel stiffness^15^. Furthermore, the formation of supramolecular DNA hydrogels is also dependent on the rigidity of the network^47^. For comparison, we evaluated the solution of a single 3WJ-Sh motif at 60 µM that did not yield a hydrogel (***Figure S18b***). At 90 µM, these values increased significantly in magnitude (150 ± 31 Pa for 3WJ-Sh, *p* = 0.048; 410 ± 130 Pa for 3WJ-Lo, *p* = 0.041; 255 ± 13 Pa for 3WJ-Lm, *p* = 0.0003) for all linkers tested, compared to 60 µM, but remained in the same order of Lo ≥ Lm > Sh, demonstrating a concentration-dependent effect as well. These moduli are comparable in magnitude to other DNA hydrogels on the order of a few hundred Pa^48^, but require much less material (∼0.2-0.3% solid content) to match and even exceed properties of similar constructs^15^. Similar to the amplitude sweep results, G’ magnitude for the 30 µM hydrogels was lower for all linkers (6 ± 2 Pa for 3WJ-Sh; 21 ± 19 Pa for 3WJ-Lo; 2 ± 1 Pa for 3WJ-Lm) (***Figure S19b***), likely due to its incomplete formation. For the Y-motif-based hydrogels, they again did not appear to form well at the motif matching concentration of 60 µM (***Figure 17a, bottom***), but the G’ was 90 ± 10 and 49 ± 33 Pa for DNA content matching concentrations of Y7-150 µM and Y21-120 µM, respectively (***Figure 17b, bottom***). Unlike the DX-based hydrogels, it has been proposed that the longer overhang for single-duplex DNA hydrogels may exceed the optimal range for gel formation^30^. Additionally, this result confirms that for the same concentration of DNA or total motif, our DX-DNAns hydrogels possess enhanced mechanical strength over Y-motif-based hydrogels and other peer designs while offering greater tunability. This result is consistent with what is known about the critical influence of the DNA structure on its physical features, such as its persistence length (∼50 nm^49^ or 150 bp) and the rigidity of the DX backbone, which is twice that of linear duplex DNA^16^.

To determine the effect of temperature on the formation of the hydrogels, we compared the rheological properties of 3WJ-Sh hydrogels measured either (1) immediately after formation or (2) following a 12-minute temperature cycle consisting of heating to 55°C (to ensure melting of only the linkers only) then cooling back to room temperature to induce reassembly (***Table S4***, ***Figure S20***). After one heat cycle, the G’ of the 3WJ-Sh hydrogels decreased significantly by ∼50% to 41 ± 10 Pa for 60 µM (*p* = 0.017) and 82 ± 14 Pa for 90 µM (*p* = 0.048) (***Table S4***). We observed an even more drastic effect of nearly 75% with Y7-based hydrogels (89.8 ± 10.0 Pa pre-heat cycle to 23.4 ± 11.2 Pa post-heat cycle) (***Figure S21***). Differences in the reduction of G’ for 3WJ-Sh and Y7 hydrogels might be explained by the higher thermal stability provided by the two linkers in the 3WJ, compared with the single linker for the Y7. Importantly, neither hydrogel recovered completely after annealing, which can be due to constrained diffusion of the motifs and incomplete disruption of the hydrogel networks.

Given that the rigidity and other physical properties of DX-tile-based nanoconstructs are dependent on their structural design, including crossover density and arm length^50–53^, we tested 3WJ-Sh constructs with different crossover densities (***Table S5***; ***Figure S22a***) and arm lengths (***Tables S6, S7***; ***Figure S1b***). For the crossover densities, the G’ was higher for hydrogels possessing one extra crossover in the middle of the arm (274 ± 45 Pa for 3WJ-Sh-X *versus* 84 ± 11 for 3WJ-Sh; ***Figure S22b***). However, the G’ was lower for constructs with an extra double crossover (14 ± 4 Pa for 3WJ-Sh-XX; ***Figure S22c***). This difference could be explained by the fine balance necessary between arm rigidity to increase stiffness and flexibility to permit hybridization with other motifs in a 3D mesh^16,47,54^. For the arm lengths, the short arm hydrogels composed of repeating sequences with either an outside (***Figure S1bi***) or inside (***Figure S1bii***) crossover did not appear to form a hydrogel (***Figures S23a*** and ***S23b***, respectively). However, when we altered the design to non-repeating sequence with an inside crossover (***Figure S1biii***), the resultant hydrogel had the same crossover point as the long arm variant (***Figure S1c***) at 102% (***Figures S23c*** and ***23d*** *left*, respectively). The arm length did affect the G’ of the hydrogels, as those formed with the non-repeating, inside-crossover variant possessed a shorter, more rigid arm and thus had a lower modulus (49.2 ± 6.7 Pa; ***Figures S23c*** *right*) than those formed with the longer, more flexible arm variant (184 ± 73 Pa; ***Figures S23d*** *right*). These values were lower and greater than the comparable regular arm analogue, respectively. This difference could be attributed to the overall flexibility of the longer arm in resisting deformation, thereby supporting its more elastic behavior.

### Arm valency in DX-tile based DNA hydrogels alters bulk mechanical properties

Next, we explored how the arm valency could impact the bulk material properties. Previous studies have shown that the valency of crosslinkers drastically influenced the physical properties of the hydrogels formed, likely due to its greater crosslinking density^55,56^. We proceeded in the same fashion as with the 3WJ by designing a six-way junction (6WJ) central core (Co) that comprised the vertex of the full motif unit (***Table S8*** and ***Figure S1d***) and was capable of displaying the same linker types (***Figure S24a***). Its arm-to-arm length was estimated at 20.7 nm and the measured arm angle at 64.5 degrees (***Figure S24b***). We confirmed proper folding of the Co and full motifs with agarose gel electrophoresis (***Figure S24c***) and AFM imaging for the individual motifs (***Figure S24d***). Additionally, we checked the melting temperatures (***Figure S24e***) of the individual Co (∼68°C) and Pl arm (∼67°C) and in combination (∼62°C and ∼68°C, respectively).

As with the 3WJ hydrogels, we tested the formation of the 6WJ hydrogels with the Sh and Lo linkers by mixing complementary motifs at equimolar concentration ranging from 7.5 to 45 µM (***Figure S25***). These concentrations were chosen to match the DNA content to that of the 3WJ hydrogels (***Figure S3***) since 60 and 90 µM motif concentrations were not possible for the 6WJ hydrogels with the additional strands per motif (16 strands for 3WJ *versus* 33 strands for 6WJ). The formation of hydrogels was not observed for low concentrations of 7.5 and 15 µM of either motif. Hydrogels were clearly formed for both motifs at 30 and 45 µM, with a clear concentration dependence on the apparent mechanical properties. SEM analysis (***Figure 4a, Figure S26a***) further confirmed hydrogel formation across the different linkers, including the Bl linker, and different concentrations with clear and extensively uniform microporous networks (***Figure S27***). While the measured Y7 (225 µM) and 3WJ-Lm (90 µM) pore sizes were similar (5.8 ± 1.5 µm *versus* 5.7 ± 0.2 µm, respectively), the 6WJ-Lm (50 µM) pore size was significantly less at 3.9 µm (*p* < 0.001). This result suggests that the motif valency extent of crosslinking seem to positively affect the reduction in hydrogel pore size. Compared to the general 3WJ pore size of ∼5-6 µm, the 6WJ pore size of < 4 µm presents another morphological feature that can be modulated along with mechanical properties.

**Figure 4.**
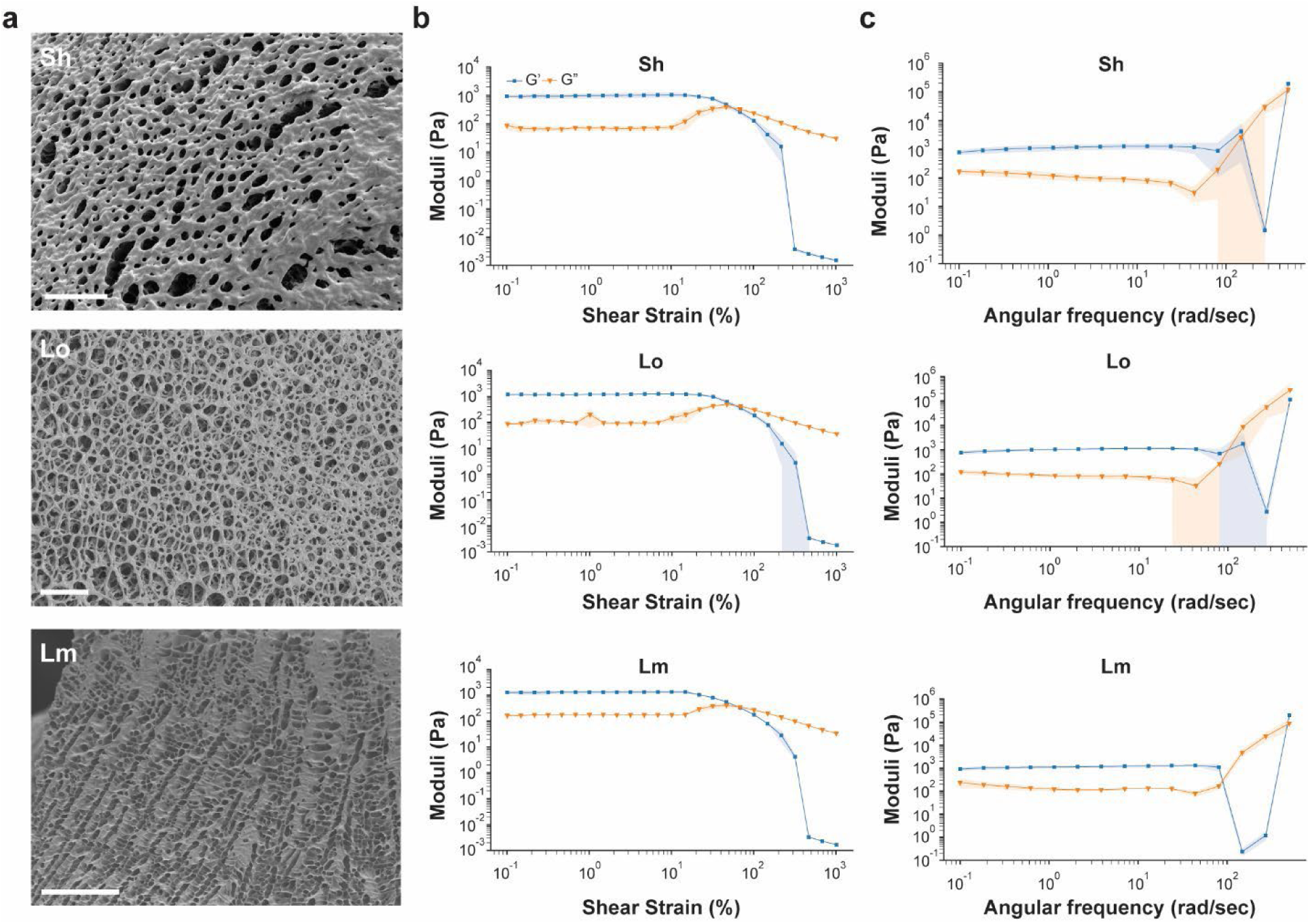
Characterization profile of the 6WJ constructs. (a) Representative SEM images (scale bar: 50 µm). (b) Amplitude sweeps. Error shadings represent standard deviation of the mean (*n* = 3 independent samples/group). (c) Frequency sweeps. Error shadings represent standard deviation of the mean (*n* = 3 independent samples/group).

We performed a series of amplitude sweeps (***Figure 4b***, ***Figure 26b***) to ascertain the LVER for the three hybridizing DX-tile motifs at two motif concentrations that were approximately comparable to some of those tested for the 3WJ (***Table S3***). For the 6WJ motif, the LVER crossover point was consistent across all three linkers and at both concentrations in ascending order of Sh (38%, 25 µM; 55%, 50 µM), Lo (40%, 60 µM; 58%, 90 µM), and Lm (52%, 60 µM; 65%, 90 µM) linkers. The difference between the concentrations was not as pronounced as the 3WJ, suggesting the influence of the valency (3WJ *versus* 6WJ) is greater than that provided by the motif concentration.

For the frequency sweeps (***Figure 4c; Figure 26c***), G’ increased in magnitude for all of the samples relative to their 3WJ counterparts. For the 25 µM hydrogels, those formed with the Lm linker had the highest G’ at 1 Hz (282 ± 25 Pa), while those with the Sh linker had the lowest G’ (119 ± 12 Pa) and those with the Lo linker had a G’ in-between the others (179 ± 46 Pa). This pattern matched that of the amplitude sweep for both hydrogel concentrations (Lm > Lo > Sh). However, this trend significantly reversed for the 50 µM hydrogels, where those formed with the Sh linker had the highest storage modulus at 1 Hz (1266 ± 230 Pa; *p* = 0.002), while those with the Lo linker had the lowest storage modulus (1095 ± 8 Pa; *p* < 0.001) and those with the Lm linker fell in-between (1201 ± 128 Pa; *p* = 0.0006). This concentration-dependent shift to Sh > Lm > Lo may result from the increased crosslinking creating a cooperative effect, as these higher moduli were all similar (*p* > 0.3), and from the shorter overhang creating greater rigidity. As Similar to the 3WJ samples, we performed a heat step and observed a decrease in the storage moduli for the 25 µM and 50 µM to 29 ± 17 Pa (*p* = 0.004) and 347 ± 29 Pa (*p* = 0.005), respectively (***Table S4, Figure S28***). The combination of a higher motif density and greater motif valency may have interfered with the crosslinking ability of complementary motifs to relocate one another after the additional heating step. A summary set of these storage moduli is displayed in ***Table 1*** (*6WJ, right*).

To further assess the precise tunability of our system, we evaluated the effect of matching the valency ratios between the 3WJ and 6WJ motifs by combining them at 2:1 (3WJ:6WJ) and verified hydrogel formation by SEM (***Figure S29a***). We tested different motif ratios of 1:6, 1:1, and 6:1 to assess their impact on rheological properties. For the amplitude sweeps (***Figure S29b***), the 1:1 and 1:6 samples had similar LVER crossover points of 39% and 38%, respectively; however, for the 6:1 sample, it was much higher at 69%. Likewise, for the frequency sweeps (***Figure S29c***), the 6:1 sample had the highest G’ modulus at ∼1 Hz (416 ± 81 Pa), while a stepwise effect was observed for the 1:6 and 1:1 ratios from 270 ± 38 Pa to 347 ± 54 Pa, respectively. Interestingly, all combination ratios produced increased stiffness relative to their single motif counterpart. This outcome could be explained by the formation of a more branched network when using multiple motifs of different valencies, thus confirming the importance of the nanoscale organization of motifs. Moreover, adjusting valency ratios between motifs not only controls the mechanical properties of our hydrogels, but can also create tailored materials amenable to patterned functionalization.

### DX-tile DNA Hydrogels demonstrate 3D printability

The viability of DX-DNAns hydrogels as extrudable bioink for 3D printing was also assessed. We first conducted rheological measurements to ensure that these new hydrogel formulations were suitable for extrusion 3D printing. For our proof of concept, we chose the 6WJ-based DNA hydrogels with the highest elastic modulus. We verified the hydrogels were shear-thinning (***Figure 5a***), confirmed the point at which their elastic behavior broke down and began to flow (***Figure 5b***), and studied if they recovered to their previous physical state (***Figure 5c***). Recoverability was of particular interest, as we had observed self-healing behavior with this material following tears and cuts (***Figure S30***).

**Figure 5.**
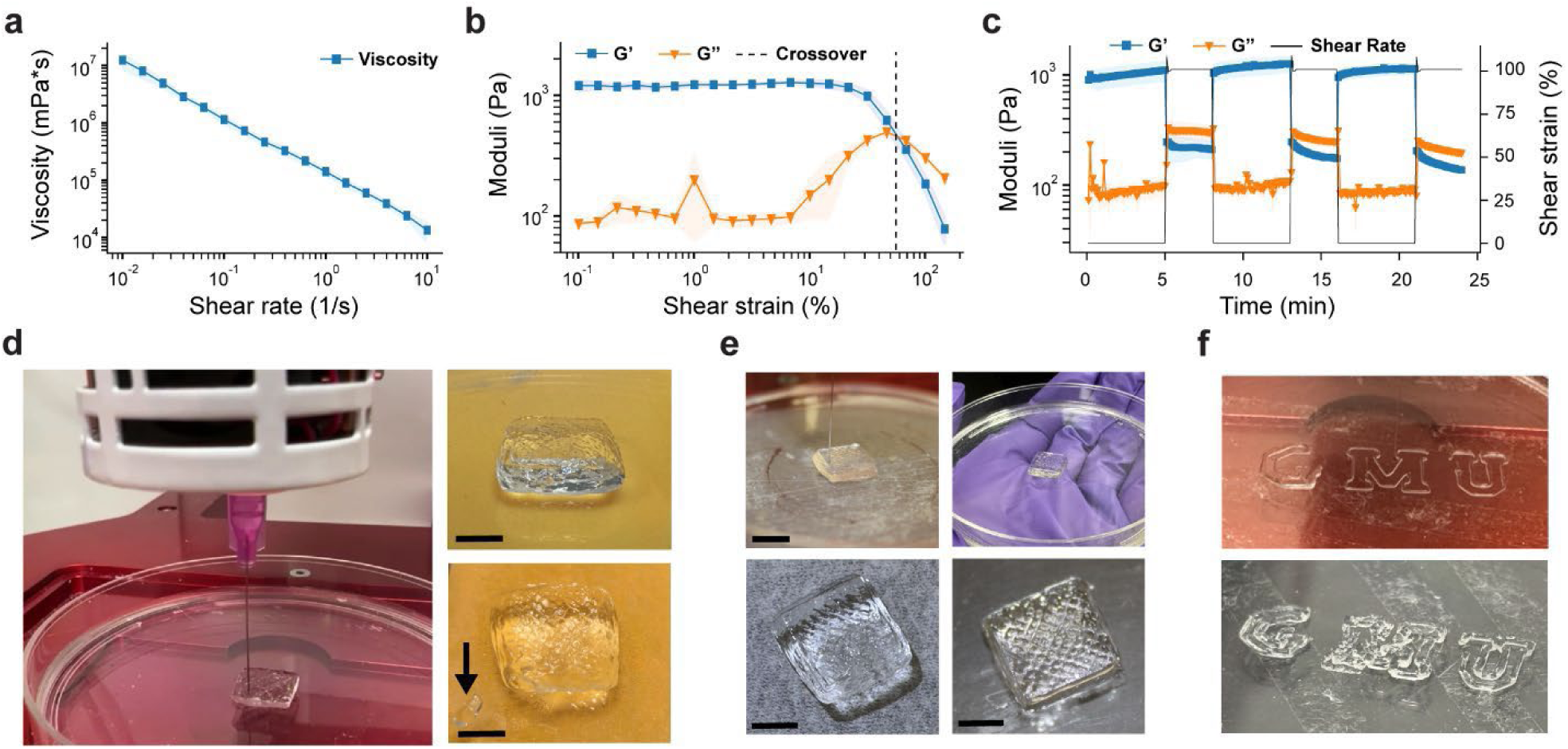
3D printing profile of 6WJ constructs. (a) Viscosity curve indicated shear thinning behavior. Error shadings represent standard deviation of the mean (*n* = 3 independent samples/group). (b) Amplitude sweep determined LVER. Error shadings represent standard deviation of the mean (*n* = 3 independent samples/group). (c) Cyclic strain testing demonstrated recoverability. Error shadings represent standard deviation of the mean (*n* = 3 independent samples/group). (d) First print with the Lm linker in process (*left*), after completion (*top right*), and after 12 hours in 4°C with a piece removed (*arrow*) to show its integrity (*bottom right*) (scale bar: 1 cm). (e) First print with the Lo linker in process (*top left*), after completion (*bottom left*), and after collection and reprinting following a brief re-annealing step at 55°C (*right top/bottom*) (scale bar: 1 cm). (f) GMU print with Lo linker in process (*top*) and after completion (*bottom*).

Once testing was complete for the basic material properties, we used an Allevi 3 extrusion printer to test the viability of 3D bioprinting the material. Various printing parameters were tested to identify a successful printing configuration: 25-gauge (G) and 30 G syringes at 30, 35, and 40 pounds per square inch (PSI) were used to create prints at 1X, 2X, and 3X the inner diameter of the needle under 6, 9, and 12 mm/s print speeds (***Figure S31***). Following successful tests, an ideal combination of parameters was identified, and a 20 x 20 x 10 mm brick print shape was modeled using CAD and sliced for printing with 130 layers. The gel was printed with a 1-inch, 30 G needle at 68 PSI and 6 mm/s speed (***Figure 5d***, *left*). After storage at 4°C for 12 hours (***Figure 5d***, *top right*), we cut a piece of the hydrogel to confirm its integrity (***Figure 5d***, *bottom right*). Next, we printed the cube again and collected the material to immediately reprint the same design with no observable reduction in print quality, thus demonstrating the self-healing properties intrinsic to these pure DNA hydrogels (***Figure 5e***)^57,58^. Finally, as another demonstration of material versatility, we printed the George Mason University (GMU) letters to exhibit the excellent print resolution and filament fidelity (***Figure 5f***).

### DX-DNAns hydrogel functionalization and stability

DNA motifs can be readily modified with a strand displacement reaction (SDR) system to enable the stimuli- and temporal-controlled release of various cargos to create smart biomaterials^59,60^. The release of cargo molecules directly attached to the motifs constituting the DNAns hydrogels affect the mechanical properties or structure of these hydrogels^61,62^. Here, we aim to demonstrate that our DX-DNAns hydrogels serve as a platform to design SDR systems capable of performing controlled release without affecting their mechanical properties or stability. Leveraging a previously published SDR circuit devised^63^, we conjugated our 3WJ and 6WJ DNA motif designs with a HEX fluorophore attached to IowaBlack quencher strand (***Figure 6a***). This circuit was designed such that adding an excess of a trigger strand with greater affinity to the modified DNA motif than the quencher strand would displace the quencher over a period of time that could be quantified by an increase in fluorescence intensity.

**Figure 6.**
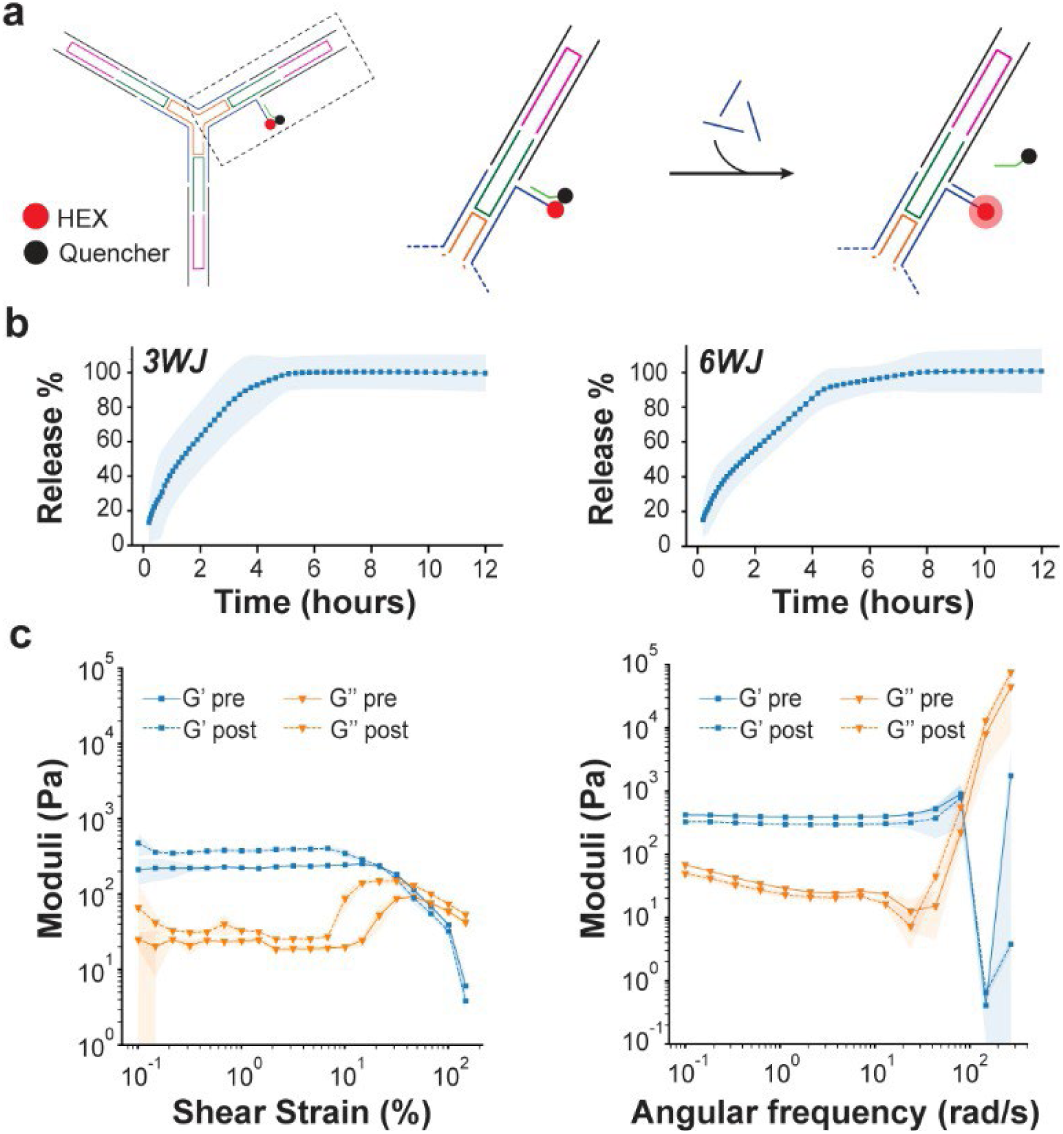
Hydrogel functionalization and stability. (a) Schematic diagram of motif modification for strand displacement release system. (b) Release curve for 3WJ (*left*) and 6WJ (*right*) hydrogels with the Sh linker at comparable concentrations of 30 µM and 60 µM, respectively. Error shadings represent standard deviation of the mean (3WJ, *n* = 3 and 6WJ, *n* = 7 independent samples/group). (c) Amplitude (*left*) and frequency sweep (*right*) of functionalized 3WJ-Sh hydrogels at 60 µM before and after cargo release with ssDNA triggers. Error shadings represent standard deviation of the mean (*n* = 3 independent samples/group).

With both constructs at comparable amounts of DNA content (3WJ, 60 μM; 6WJ, 30 μM), we observed a similar cargo release between these two motif types, with only minor and non-significant differences in their kinetic profiles (3WJ: 0.54 ± 0.25 hr^-1^ *versus* 6WJ: 0.44 ± 0.26 hr^-1^; *p* = 0.69) and 50% release time (3WJ 1.29 ± 0.79 hr *vs* 6WJ 1.57 ± 1.41 hr; *p* = 0.67) (***Figure 6b***). This small difference in their release could be attributed to the smaller pore sizes with the 6WJ construct. However, the similarity of their release profiles is notable, given the significant differences in their individual mechanical properties. The complete set of results is summarized in ***Table S9***.

To verify that functionalization did not disrupt the mechanical properties of the hydrogels, we performed rheology tests on the 3WJ hydrogels loaded with the SDR system before strand triggering and after release. As determined by amplitude (***Figure 6c* *left***) and frequency sweeps (***Figure 6c* *right***), the rheological profile remains largely intact after completely triggering the strand release in the hydrogel. The LVER crossover point shifted between pre- and post-triggering from 32% to 60%, respectively, and G’ at ∼1 Hz changed from 391 ± 11.26 to 298 ± 15.76 Pa after total strand release. The ∼24% decrease in G’ is far less than previously reported for similar systems^47^ and is consistent with the stability of the mechanical properties for unmodified hydrogels at different timepoints after preparation (***Figure S32***). For one gel (3WJ-Sh, 60 µM) analyzed over Days 1, 3, and 5 post-folding, the storage modulus at ∼1 Hz was 202 ± 33 Pa, 203 ± 32, and 169 ± 31 Pa, respectively, which may represent spontaneous and dynamic reorganization of the hydrogel over time or slow degradation overall.

### Conclusion

DNA nanotechnology allows unprecedented control over molecular and nanostructural design elements that enable precise functionalization. However, the development of macroscale materials made purely from DNA has been limited to duplex-based DNAns motifs and long hybridizing DNA, with little exploration into more complex structures and design features that can tune the physical properties of these materials. DNA nanostructures, for instance, are limited to a few hundred nanometers in size, depending on scaffold length, and remain relatively costly to produce, which fundamentally hinders the production of large-scale materials^64,65^. Furthermore, it is unclear to what extent these nanostructures can be linked to form macroscale materials, or whether the structures, or their crosslinks, will remain stable.

We explored the impact of controlling the critical physical parameters of our DX-DNAns hydrogels, from linker and arm design to arm valency, which affect the bulk mechanical properties, while keeping the structure intact and motif concentrations identical. Our results demonstrated that the nanoscale flexibility afforded by these DX-tile motifs can produce programmable, self-organizing macroscale hydrogels with highly tunable mechanical properties involving material stiffness from less than 30 Pa to more than 1.2 kPa. Moreover, DX-DNAns-based hydrogels solve the limitations of previous iterations of pure DNA hydrogels, offering a unique medium to study soft robotics and polymeric materials with design control and efficient synthesis. In addition, given its similarity with 3D DNA nanostructures, this technology could enable nanoscale organization and manipulation of various biomolecules for to direct cell behavior and regulate lipid membranes, deliver drugs and targeted therapies, determine biophysical interactions, and analyze molecular structures^66–68^. From the nano- to macro-scale, this unique strategy presents a desirable candidate for the rational design of hydrogels with prescribed mechanical properties that can be leveraged for biological applications as varied as studying viscoelastic influences on mechanobiology^33^ to designing scaffolds for soft tissue engineering, such as adipose and nerve tissues^54^.

## Methods

### Materials

All DNA oligonucleotides, modified (i.e., Cy3, IowaBlack, HEX) and non-modified, were purchased from IDT and directly used without further purification. All sequences used are listed in the Supplementary Tables S1-S2; S5-S8. The stock folding buffer (10X) used to assemble the motifs is composed of TAE buffer (40 mM Tris, 20 mM acetic acid, 2 mM ethylenediaminetetraacetic acid) plus 12 mM magnesium chloride hexahydrate (MgCl_2_) at pH 8.0. The folding buffer is used at 1X, with the remaining volume being nuclease-free water (E476; VWR Life Science). For gel electrophoresis, high-melt agarose (IB70070; IBI Scientific) was used at 1.2% with 1X TAE (pH 8.0) and pre-stained with 0.5 μg/mL ethidium bromide (15585-011; Invitrogen) while samples were loaded with 1X Gel Loading Dye Purple (B7025S; New England BioLabs). Array samples were artificially colored with printer linker (X001OPYOXP; Aomya). qPCR samples were stained with 1X SYBR™ Green I Nucleic Acid Gel Stain (50513; Lonza) and diluted in folding buffer. SEM samples were flash-frozen in liquid nitrogen (Roberts Oxygen) and mounted with carbon tape (77825-12; Electron Microscopy Sciences) on aluminum pedestals (RS-MN-10-005112-50; RaveScientific). Positive displacement pipettes (Gilson) were used for handling hydrogels.

### Design of the DX-tile DNA motifs

The DX-tile based DNA motifs were designed with the Tiamat 2 program^69^. The strand sequences designed for each construction are available in the Supplementary Tables S1 to S3. To ensure that a designed strand sequence would only bind to its desired complementary sequence and avoid non-specific binding, we used NCBI’s Nucleotide BLAST^70,71^.

### oxDNA modeling of the DNA motifs and visualization of the PDB models

All DNA motifs created in Tiamat 2 were converted into PDB using the TacoxDNA tools suite^72^. The University of California San Francisco’s Chimera 1.16^73^ was used to visualize the 3D molecular structure of the motif and estimate distances and angles.

### Folding the DNA motifs

The DNA motifs were folded in a one pot reaction with all strands added at equimolar concentration or at indicated molar ratios for strands used multiple times in a motif (***Tables S1-S3***). The oligonucleotide mix was prepared in our 1X folding buffer and annealed from 90 to 4°C overnight in a Biorad T100 thermocycler. Following folding, the nanostructures were stored at 4°C without further purification prior to characterization or gel formation. The concentration of the DNA motifs was measured using a Nanodrop One. Folded DNA motifs were run on 1.5% agarose gel pre-stained with ethidium bromide at 100 V for 30 minutes and imaged with an Azure c150 blue transilluminator.

### FRET assay

Folded structures were validated via FRET assay performed with a Cy3/IowaBlack dye quencher pair. Cy3 dye was excited at 555 nm and the maximum emission was monitored at 570 nm. Samples were prepared at a volume of 40 µL into a 384-well black flat transparent bottom plate.

### Gel assembly

All hydrogels were assembled by mixing equal parts of one motif (A motif) and its complementary form (B motif) in 1X folding buffer. The only exception were the blunt-end DNA motifs, which did not require a complementary moiety as they did not have linkers. The hydrogel concentrations reported refer to the total motif concentration.

### qPCR melting curves

Melting curves were acquired using The Q instrument from Quantabio. A total of 100 µL for each sample was pipetted evenly into a strip of four loading tubes (1.25 µL of 1X SYBR Green I dye, 12.5 µL of motif, 11.25 µL of folding buffer per tube). Light exposure was minimized to avoid bleaching of the dye. The tubes came preloaded from the supplier with silicone oil to prevent evaporation, condensation, and the need for a heated lid. The samples were melted from 40.0°C to 95.0°C at 0.05°C/s, for two cycles. Melting curves were extracted from the qPCR cycler as Excel files, and the melting temperature was determined by taking the average of the fluorescent intensity for the quadruplicate samples. Data was then normalized against the peak value.

### Atomic Force Microscopy evaluation

AFM images were captured with a Bruker Dimension Icon + ScanAsyst. For imaging the DNA hydrogels, 60 µM hydrogel samples were layered onto freshly cleaved mica surfaces (Electron Microscopy Sciences) and left in a vacuum chamber overnight to dry. The samples were then imaged in ScanAsyst mode with a SCANASYST-AIR-HPI probe (Bruker). For DNA motif imaging, 20 μL of ∼1-5 nM folded DNA motifs in folding buffer were deposited onto freshly cleaved mica and incubated for 10 minutes, followed by several gentle folding buffer washes. The motifs were immediately imaged in folding buffer supplemented with 10 mM NiCl_2_, using ScanAsyst mode and a SNL-10 probe (Bruker). All AFM images were processed using Gwyddion^74^.

### Dynamic Light Scattering measurements

Hydrodynamic diameters of the folded 3WJ motifs were measured via DLS using a Malvern Zetasizer Nano ZS instrument. Samples were diluted with 1X folding buffer to a final concentration of 1 µM in a volume of 60 µL and loaded into disposable plastic cuvettes. Size measurements were taken with 173° backscatter.

### Scanning Electron Microscopy evaluation

SEM was used to confirm hydrogel formation with the JEOL JSM-7200F instrument and PC-SEM software. Formed hydrogels (40 µL) were directly cast into 3D-printed resin rings (12-mm diameter) that had been fixed onto brass pedestals with carbon tape and then freeze-dried with at least 10 minutes of liquid nitrogen and 24 hours of lyophilization. Samples were sputter-coated with gold using a Denton Desk V instrument for 30 seconds at 2 mA and imaged with 1-3 kV of accelerating voltage.

### Pore Size analysis

Fully formed pores (50 count) were selected across 4-5 in-plane SEM images (∼400-500x magnification) and measured with ImageJ for each of the various hydrogels. The scale was set using the “Set Scale” function and the rectangle tool to match that of the SEM image, and the pores were measured using the straight-line tool to find their X and Y lengths. The X measurement spanned the shorter distance and bisected the pore into equal halves, while the Y measurement was perpendicular to this initial measurement and spanned the longer distance. The pore size measurements were collated into Excel for analysis by hydrogel design, including average cell size and standard deviation.

### Rheology measurements

Nanostructures were folded at indicated molar concentrations in 1x folding buffer. Identical complementary structures were mixed on the Anton Paar MCR302 rheometer plate via pipette prior to rheology measurements. Rheology was conducted with a parallel plate setup (non-sandblasted Peltier plate) using the PP08/S measuring system (97676; Anton Paar), and measurements were taken at a strain of 1%. Measurements were taken after mixing and then again after a heating and cooling cycle. The heating cycle increased linearly from 25 to 55°C over 5 minutes with a 2-minute hold at 55°C followed by a cooling ramp to 25°C over the course of 5 minutes. The hydrogel was allowed to set for 5 minutes at 25°C before the second set of rheology measurements were taken.

### Printing tests

Prints were conducted at an ambient temperature ranging from 20-23°C using Allevi 3 extrusion 3D printer (Allevi, Inc.). Initial line tests were performed to identify ideal printing parameters (***Figure S29***); the test indicated ideal parameters of 68 pounds PSI pressure with a 1-inch, 30-gauge nozzle. Subsequent tests were performed with rectilinear infill, printing a cube of 2 cm x 2 cm x 1 cm, as well as “GMU” block letters with one infill per layer. Rectilinear cube prints were inspected for cohesion and integrity to print vertically without deformation. Subsequently, the material was recollected, heated, recentrifuged, and reprinted (**Figure 5e**). “GMU” block letters were printed to identify the material’s ability to support its own weight with only a single layer infill per Z stack, as well as the ability to manipulate in 90° corners and sweeping curves (**Figure 5f**).

### Implementation of a DNA strand release system in the hydrogel

The functionalization of the DNA hydrogels was assessed by quantifying the release of a quencher from the HEX-modified DNA motifs (1μM) and measuring the increase in fluorescence intensity for each fluorophore after the addition of a trigger strand (2:1 trigger:motif) at 40 °C using the ‘Q’ qPCR machine (Quantabio). The protocol was set to run for 60 cycles with 120 s of cycling time after a 5-min hold period, wherein the cycling time increased 20 s per cycle (e.g., 140 s of cycling time for cycle two and 160 s for cycle three) for a total run time of ∼12 h. The excitation for HEX was 540 nm, and the emission was recorded at 570 nm. The release was normalized against the positive control, which was the DNA motif without the quencher, and all the values are reported as the mean ± standard deviation.

### Hydrogel mechanical stability assay

The mechanical stability of the 3WJ-Sh hydrogels at 60 µM was determined with frequency sweeps at the indicated post-assembly timepoints (0, 4, 5 days) as previously described in the Rheology Measurements section. Between measurements, hydrogels were stored at 4°C.

### Statistical analysis and reproducibility

All tests were performed in triplicates and all values were reported as the mean ± standard deviation, unless otherwise indicated. For the pore size measurements and rheology analyses, two-tailed student t-tests were conducted assuming two samples of equal variance. *P*-values were reported with specified thresholds for significance or non-significance. All statistical analyses were conducted with Excel.

## Acknowledgments

This work was supported by The Assistant Secretary of Defense for Health Affairs endorsed by the Department of Defense, in the amount of $317,780.00 through the Peer Reviewed Medical Research Program under Award Number HT9425-23-1-0037. Opinions, interpretations, conclusions, and recommendations are those of the author(s) and are not necessarily endorsed by The Assistant Secretary of Defense for Health Affairs endorsed by the Department of Defense. A Cosmos Scholars Grant from the Cosmos Club Foundation was awarded to D.V.S. to support the purchase of experimental supplies. Additionally, the bioprinting contributions were supported by the Center for Rehabilitation Sciences Research through the Department of Physical Medicine & Rehabilitation at the Uniformed Services University of the Health Sciences in Bethesda, MD (Award #HU00012320007). The opinions and assertions expressed herein are those of the authors and do not necessarily reflect the official policy or position of the Uniformed Services University, Department of Defense, or Henry M. Jackson Foundation for the Advancement of Military Medicine, Inc. Additionally, We would like to thank Joshua Bush for his contributions to the early stage of this work.

## Contributions

Conceptualization, R.V.; methodology, D.V.S., S.J., R.V.; investigation, D.V.S., A.B.C., E.O.T., K.A.H., P.M.T., J.B., R.C.S., C.R.F., H.G.M., C.H., R.V.; original draft preparation, D.V.S., R.V.; writing: review and editing, all authors; supervision, X.Y., S.J., R.V.; funding acquisition, D.V.S., X.Y., S.J., R.V. All authors have read and agreed to the published version of the manuscript.

## Supplementary information

Supplementary Figs. 1–32 and Tables 1–9.

