## Supplementary Information for "Designer DX-tile DNAns hydrogels"

**Table S1. DNA sequences for double-crossover (DX)-tile three-way junction (3WJ) motif with regular arm and blunt (Bl), short (Sh), long (Lo), or long mismatch (Lm) linkers, including variants with one (X) and two (XX) additional crossovers.** Regions highlighted in orange represents hybridizing sticky ends.

| Component (xMolar Ratio) | | Strand | Sequence |
|---|---|---|---|
| **Core (x1)** | | 1 | GGAAGTCTAGACAGTAGCGGCTTCCGCAAGATTTCACCAGGG |
| | | 2 | GGAAGTCTAGCGATATAAATTTGTCGGGACTTTTCACCAGGG |
| | | 3 | GGAAGTCTAGCGTAAGTCTCGGGGGTAGGAATTTCACCAGGG |
| | | 4 | CGGAATTTGCCGCTACTGTAGTCCCGACATTTAATTTATATCGTTCCTACC CCTTTCGAGACTTACGTCTTG |
| **Arm (x3)** | | 5(X) | AGGCACGGACCCCTGGTGAAACTAGACTTCCCCCTACATTTT |
| | | 5XX | ACCCCTGGTGAAACTAGACTTCCCCCTACATTTTAGGCACGG |
| | | 6 | AATCCCAGCCCGCTATGGACGTTGGTCTACGCGACCCCTCCG |
| | | 6X | GACCCCTCCGAATCCCAGCCCGCTATGGACGTTGGTCTACGC |
| | | 6XX | TATGGACGTTGGTCTACGCGACCCCTCCGAATCCCAGCCCGC |
| **Linkers (x3)** | **Bl** | 7 | CGTCCATAGCGGGCTGGGATTAAAATGTAGGG |
| | | 8 | GTCCGTGCCTCGGAGGGGTCGCGTAGACCAA |
| | **Sh** | 7 | AGACATCCGTCCATAGCGGGCTGGGATTAAAATGTAGGG |
| | | 8 | GTCCGTGCCTCGGAGGGGTCGCGTAGACCAACCTGCAC |
| | | 7' | GATGTCTCGTCCATAGCGGGCTGGGATTAAAATGTAGGG |
| | | 8' | GTCCGTGCCTCGGAGGGGTCGCGTAGACCAAGTGCAGG |
| | **Lo** | 7 | GAATTAATATATATAGACATCCGTCCATAGCGGGCTGGGATTAAAATGTA GGG |
| | | 8 | GTCCGTGCCTCGGAGGGGTCGCGTAGACCAACCTGCACTTTGTTTGGCTA GG |
| | | 7' | GATGTCTATATATATTAATTCCGTCCATAGCGGGCTGGGATTAAAATGTA GGG |
| | | 8' | GTCCGTGCCTCGGAGGGGTCGCGTAGACCAACCTAGCCAAACAAAGTGC AGG |
| | **Lm** | 7 | GAATTAATTTTTTTAGACATCCGTCCATAGCGGGCTGGGATTAAAATGTA GGG |
| | | 8 | GTCCGTGCCTCGGAGGGGTCGCGTAGACCAACCTGCACTTTTTTTGGCTAG G |
| | | 7' | GATGTCTTTTTTTTTTAATTCCGTCCATAGCGGGCTGGGATTAAAATGTAG GG |
| | | 8' | GTCCGTGCCTCGGAGGGGTCGCGTAGACCAACCTAGCCTTTTTTTGTGCAG G |

**Table S2. Y motif sequences.** Regions highlighted in orange represents hybridizing sticky ends.

| Y Motifs | | |
|---|---|---|
| **Component (xMolar Ratio)** | **Strand** | **Sequence** |
| **Y7** | | |
| **A Motif (x1)** | A1 | CAGAGCTGTCCGTGAGTGAGGCAGGATGGACGCCCGCGCTGATCCGTGA |
| | A2 | CAGAGCTTCACGGATCAGCGCGGGCGTCTAACTTCCTGGCAGGCCGACT |
| | A3 | CAGAGCTAGTCGGCCTGCCAGGAAGTTACATCCTGCCTCACTCACGGAC |
| **B Motif (x1)** | B1 | AGCTCTGGTCCGTGAGTGAGGCAGGATGGACGCCCGCGCTGATCCGTGA |
| | B2 | AGCTCTGTCACGGATCAGCGCGGGCGTCTAACTTCCTGGCAGGCCGACT |
| | B3 | AGCTCTGAGTCGGCCTGCCAGGAAGTTACATCCTGCCTCACTCACGGAC |
| **Y21** | | |
| **A Motif (x1)** | A1 | GAATTAATATATATAGACATCGTCCGTGAGTGAGGCAGGATGGACGCCCGCGCTGATCCGTGA |
| | A2 | GAATTAATATATATAGACATCTCACGGATCAGCGCGGGCGTCTAACTTCCTGGCAGGCCGACT |
| | A3 | GAATTAATATATATAGACATCAGTCGGCCTGCCAGGAAGTTACATCCTGCCTCACTCACGGAC |
| **B Motif (x1)** | B1 | GATGTCTATATATATTAATTCGTCCGTGAGTGAGGCAGGATGGACGCCCGCGCTGATCCGTGA |
| | B2 | GATGTCTATATATATTAATTCTCACGGATCAGCGCGGGCGTCTAACTTCCTGGCAGGCCGACT |
| | B3 | GATGTCTATATATATTAATTCAGTCGGCCTGCCAGGAAGTTACATCCTGCCTCACTCACGGAC |

**Table S3. Approximate comparison of total DNA ratios used for each tested 3WJ and 6WJ motif concentration and their corresponding Y motif equivalent.** Total DNA concentrations were calculated by adding the molecular weights of all the motif components, including core and arm sub-units, and deriving them from the given motif concentrations. Among these tested motif concentrations, cells shaded in orange represent motif concentrations that failed to form gels, while yellow shading represents partial gel formation and green represents definitive gel formation.

| ***Total DNA concentration (mg/mL)*** | ***Y7*** | ***Y21*** | ***3WJ-Lo/Lm*** | ***6WJ-Lo/Lm*** |
|---|---|---|---|---|
| | **Motif Concentration (µM)** | | | |
| ***3.6*** | 69 | 54 | 15 | N/A |
| ***7.1*** | 139 | 108 | 30 | 15 |
| ***14.2*** | 278 | 216 | 60 | 30 |
| ***21.3*** | 417 | 324 | 90 | 45 |

**Table S4. Comparison of storage moduli at one hertz after heat cycling for 3WJ-Sh and 6WJ-Sh hydrogels.** For 3WJ (60 µM, 90 µM) and 6WJ (25 µM, 50 µM) hydrogels, baseline storage modulus is compared to that following the heat cycle protocol. (Cond.: Experimental conditions)

| **Name** | **3WJ** | | | | **6WJ** | | | |
|---|---|---|---|---|---|---|---|---|
| **[motifs]** | **60 µM** | | **90 µM** | | **25 µM** | | **50 µM** | |
| Cond. | **Base** | **Heat** | **Base** | **Heat** | **Base** | **Heat** | **Base** | **Heat** |
| G’ | 84 +/- 11 | 41 +/- 10 | 150 +/- 31 | 82 +/- 14 | 118 +/- 12 | 29 +/- 17 | 1266 +/- 230 | 347 +/- 29 |

**Table S5. Modified DNA sequences for DX-tile 3WJ motif with either one (X) or two (XX) additional crossovers in the arm.** These sequences replace the corresponding strands in the regular 3WJ motifs.

| Component (xMolar Ratio) | Strand | Sequence |
|---|---|---|
| **X Arm (x3)** | 6 | GACCCCTCCGAATCCCAGCCCGCTATGGACGTTGGTCTACGC |
| **XX Arm (x3)** | 5 | ACCCCTGGTGAAACTAGACTTCCCCCTACATTTTAGGCACGG |
| | 6 | TATGGACGTTGGTCTACGCGACCCCTCCGAATCCCAGCCCGC |

**Table S6. DNA sequences for DX-tile 3WJ motifs with short arm variants and 7-bp overhang.** Crossover (X) location indicated by name and represented in corresponding schematic: Repeating X-Out (***Figure S1bi***) or X-In (***Figure S1bii***) and Non-Repeating X-In (***Figure S1biii***). Regions highlighted in orange represents hybridizing sticky ends.

| Component (xMolar Ratio) | | Strand | Sequence |
|---|---|---|---|
| Repeating Arms | Shared Strands (x1) | 1 | AGACATCCGTCCATAGCGGGCTGGGATTCGTAAGTCTCGGGGGTAGGAA<br>CGGAGGGGTCGCGTAGACCAACCTGCAC |
| | | 1’ | GATGTCTCGTCCATAGCGGGCTGGGATTCGTAAGTCTCGGGGGTAGGAA<br>CGGAGGGGTCGCGTAGACCAAGTGCAGG |
| | | 2 | AGACATCCGTCCATAGCGGGCTGGGATTCGATATAAATTTGTCGGGACT<br>CGGAGGGGTCGCGTAGACCAACCTGCAC |
| | | 2’ | GATGTCTCGTCCATAGCGGGCTGGGATTCGATATAAATTTGTCGGGACTC<br>GGAGGGGTCGCGTAGACCAAGTGCAGG |
| | | 3 | AGACATCCGTCCATAGCGGGCTGGGATTACAGTAGCGGCTTCCGCAAGA<br>CGGAGGGGTCGCGTAGACCAACCTGCAC |
| | | 3’ | GATGTCTCGTCCATAGCGGGCTGGGATTACAGTAGCGGCTTCCGCAAGA<br>CGGAGGGGTCGCGTAGACCAAGTGCAGG |
| | | 4 | CGGAATTTGCCGCTACTGTAGTCCCGACATTTAATTTATATCGTTCCTAC<br>CCCTTTCGAGACTTACGTCTTG |
| | X-Out Strand (x3) | 5 | AATCCCAGCCCGCTATGGACGTTGGTCTACGCGACCCCTCCG |
| | X-In Strand (x3) | 5 | TTGGTCTACGCGACCCCTCCGAATCCCAGCCCGCTATGGACG |
| Non-Repeating Arms | X-In Strands (x1) | 1 | AGACATCCGCACATTGTTGGAATCAGGCCGTAAGTCTCGGGGGTAGGAA<br>TTTCACCAGGGGTCCGTGCCTCCTGCAC |
| | | 1’ | GATGTCTCGCACATTGTTGGAATCAGGCCGTAAGTCTCGGGGGTAGGAA<br>TTTCACCAGGGGTCCGTGCCTGTGCAGG |
| | | 2 | AGACATCTTAGAAGCGGTGACTTGACCTACAGTAGCGGCTTCCGCAAGA<br>CCAGTAAGTCGTGGCGCCATCCCTGCAC |
| | | 2’ | GATGTCTTTAGAAGCGGTGACTTGACCTACAGTAGCGGCTTCCGCAAGA<br>CCAGTAAGTCGTGGCGCCATCGTGCAGG |
| | | 3 | AGACATCAAAATGTAGGGGGAAGTCTAGCGATATAAATTTGTCGGGACT<br>CGTATTGCCCAGTGTGCGGTACCTGCAC |
| | | 3’ | GATGTCTAAAATGTAGGGGGAAGTCTAGCGATATAAATTTGTCGGGACT<br>CGTATTGCCCAGTGTGCGGTAGTGCAGG |
| | | 4 | TCTTGCGGAATTTGCCGCTACTGTAGTCCCGACATTTAATTTATATCGTTC<br>CTACCCCTTTCGAGACTTACG |
| | | 5 | GATGGCGCCACGACTTACTGGGCCTGATTCCAACAATGTGCG |
| | | 6 | TACCGCACACTGGGCAATACGAGGTCAAGTCACCGCTTCTAA |
| | | 7 | AGGCACGGACCCCTGGTGAAACTAGACTTCCCCCTACATTTT |

**Table S7. DNA sequences for DX-tile 3WJ motifs with long arm and Sh linker.** Regions highlighted in orange represents hybridizing sticky ends.

| Component (xMolar Ratio) | Strand | Sequence |
|---|---|---|
| Core (x1) | 1 | GGAAGTCTAGACAGTAGCGGCTTCCGCAAGATTTCACCAGGG |
| | 2 | GGAAGTCTAGCGATATAAATTTGTCGGGACTTTTCACCAGGG |
| | 3 | GGAAGTCTAGCGTAAGTCTCGGGGGTAGGAATTTCACCAGGG |
| | 4 | CGGAATTTGCCGCTACTGTAGTCCCGACATTTAATTTATATCGTTCCTACC CCTTTCGAGACTTACGTCTTG |
| Long Arm (x3) | 5 | AGTTGGCACACCCTGGTGAAACTAGACTTCCAGACGGTGCGA |
| | 6 | GCCGATGTCCGCGCCAGCTGGGAAGCAAACTGGGGAACGGAT |
| | 7 | TCACTACGTCGACCCCGCCTCTGAGGTGCCCCGCGGCCACAC |
| Sh Linker (x3) | 8 | TGTGCCAACTCCAGCTGGCGCGGACATCGGCGTGTGGCCGCGGGGCACC TCACCTGCAC |
| | 9 | AGACATCGAGGCGGGGTCGACGTAGTGAATCCGTTCCCCAGTTTGCTTCT CGCACCGTCT |
| | 8’ | TGTGCCAACTCCAGCTGGCGCGGACATCGGCGTGTGGCCGCGGGGCACC TCAGT GCAGG |
| | 9’ | GATGTCTGAGGCGGGGTCGACGTAGTGAATCCGTTCCCCAGTTTGCTTCT CGCACCGTCT |

**Table S8. DX-tile six-way junction (6WJ) sequences.** Linker sequences are the same as for the 3WJ. Regions highlighted in orange represents hybridizing sticky ends.

| Component (xMolar Ratio) | | Strand | Sequence |
|---|---|---|---|
| Core (x1) | | 1 | AGCTATCCCCTTTTTCGAGACTTACGCCTCTGGAGTTTTTTCTTGCCAGGCA |
| | | 2 | GGAAGTCTAGCGTAAGTCTCGGGGGATAGCTTTTCACCAGGG |
| | | 3 | GGAAGTCTAGGACGCCAGGAGTATATATACGTTTCACCAGGG |
| | | 4 | TTCCTACCCCTTTTTAAACTACTGTACGTATATATATTTTTCTCCTGGCGTC |
| | | 5 | GGAAGTCTAGTGCCTGGCAAGACTCCAGAGGTTTCACCAGGG |
| | | 6 | TCTTGCGGAATTTTTGCCGCTACTGTAGTCACGACATTTTTAATTTATATCG |
| | | 7 | GGAAGTCTAGACAGTAGCGGCTTCCGCAAGATTTCACCAGGG |
| | | 8 | GGAAGTCTAGCGATATAAATTTGTCGTGACTTTTCACCAGGG |
| | | 9 | GGAAGTCTAGTACAGTAGTTTGGGGTAGGAATTTCACCAGGG |
| Arm (x6) | | 10 | AGGCACGGACCCCTGGTGAAACTAGACTTCCCCCTACATTTT |
| | | 11X | GACCCCTCCGAATCCCAGCCCGCTATGGACGTTGGTCTACGC |
| | | 11ON | AATCCCAGCCCGCTATGGACGTTGGTCTACGCGACCCCTCCG |
| Linkers (x6) | Bl | 12 | CGTCCATAGCGGGCTGGGATTAAAATGTAGGG |
| | | 13 | GTCCGTGCCTCGGAGGGGTCGCGTAGACCAA |
| | Sh | 12 | AGACATCCGTCCATAGCGGGCTGGGATTAAAATGTAGGG |
| | | 13 | GTCCGTGCCTCGGAGGGGTCGCGTAGACCAACCTGCAC |
| | | 12' | GATGTCTCGTCCATAGCGGGCTGGGATTAAAATGTAGGG |
| | | 13' | GTCCGTGCCTCGGAGGGGTCGCGTAGACCAAGTGCAGG |
| | Lo | 12 | GAATTAATATATATAGACATCCGTCCATAGCGGGCTGGGATTAAAATGTAGG |
| | | 13 | GTCCGTGCCTCGGAGGGGTCGCGTAGACCAACCTGCACTTTGTTTGGCTAGG |
| | | 12' | GATGTCTATATATATTAATTCCGTCCATAGCGGGCTGGGATTAAAATGTAGGG |
| | | 13' | GTCCGTGCCTCGGAGGGGTCGCGTAGACCAACCTAGCCAAACAAAGTGCAGG |
| | Lm | 12 | GAATTAATTTTTTTAGACATCCGTCCATAGCGGGCTGGGATTAAAATGTAGGG |
| | | 13 | GTCCGTGCCTCGGAGGGGTCGCGTAGACCAACCTGCACTTTTTTTGGCTAGG |
| | | 12' | GATGTCTTTTTTTTTAATTCCGTCCATAGCGGGCTGGGATTAAAATGTAGGG |
| | | 13' | GTCCGTGCCTCGGAGGGGTCGCGTAGACCAACCTAGCCTTTTTTTGTGCAGG |

**Table S9. Functional strand release profile in 3WJ-Sh and 6WJ-Sh hydrogels.** Summary results of the functional strand release system with hexachloro-fluorescein (HEX) fluorophore and IowaBlack quencher as cargo model, showing complete saturation (*Plateau*), release rate (*k, $hr^{-1}$ and $min^{-1}$*), $r^2$ value of linear regression, and half-time course (*hr*) (3WJ n = 3 and 6WJ n = 7 independent samples/group). 3WJ and 6WJ hydrogels were made with the Sh linker at comparable concentrations of 60 µM and 30 µM, respectively.

| **Functional Strand Release Profile** | | | | | |
|---|---|---|---|---|---|
| **3WJ-Sh-60µM** | | | **6WJ-Sh-30µM** | | |
| **Plateau** | 1022.5 ± 76.9 | nM | **Plateau** | 1021.9 ± 256.3 | nM |
| **k** | 0.54 ± 0.25 | $hr^{-1}$ | **k** | 0.44 ± 0.26 | $hr^{-1}$ |
| | 32.15 | $min^{-1}$ | | 26.47 | $min^{-1}$ |
| **$r^2$** | 0.99 | --- | **$r^2$** | 0.99 | --- |
| **Half-Time** | 1.29 ± 0.79 | hr | **Half-Time** | 1.57 ± 1.41 | hr |

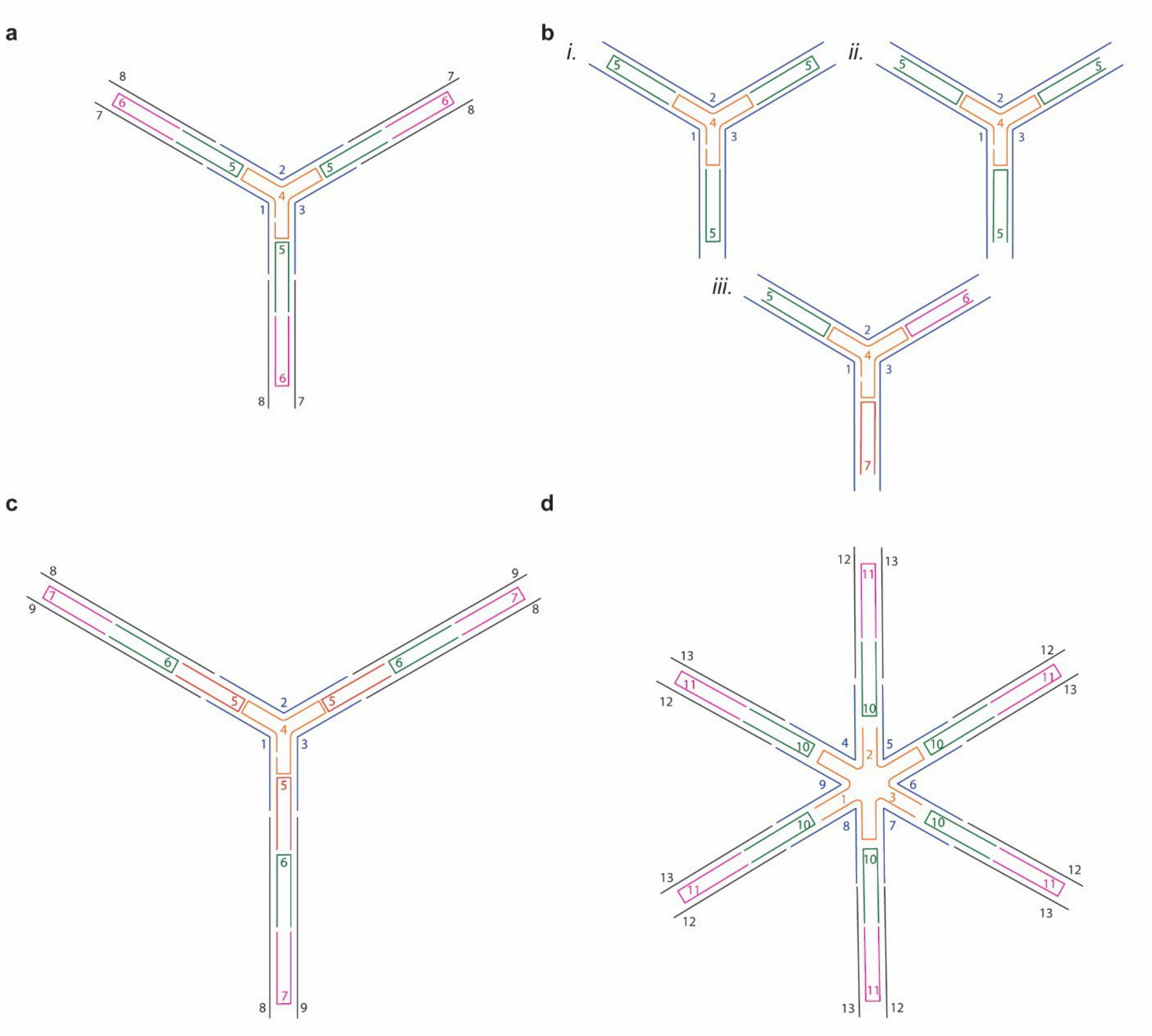


**Figure S1. Strand numbering schematic for the three DX-tile 3WJ motifs and the DX-tile 6WJ motif used in this study, all shown here with the Sh linker**. **a)** 3WJ with regular arms (53 bp). **b)** three variations of the 3WJ with short arms (32 bp) for crossover (X) locations on the repeating internal arm strand placed outside (*X-out, i*) or inside (*X-in, ii*) and the non-repeating internal arm strand placed inside (*X-in, iii*). **c)** 3WJ with long arms (74 bp). **d)** 6WJ with regular arms (53 bp).

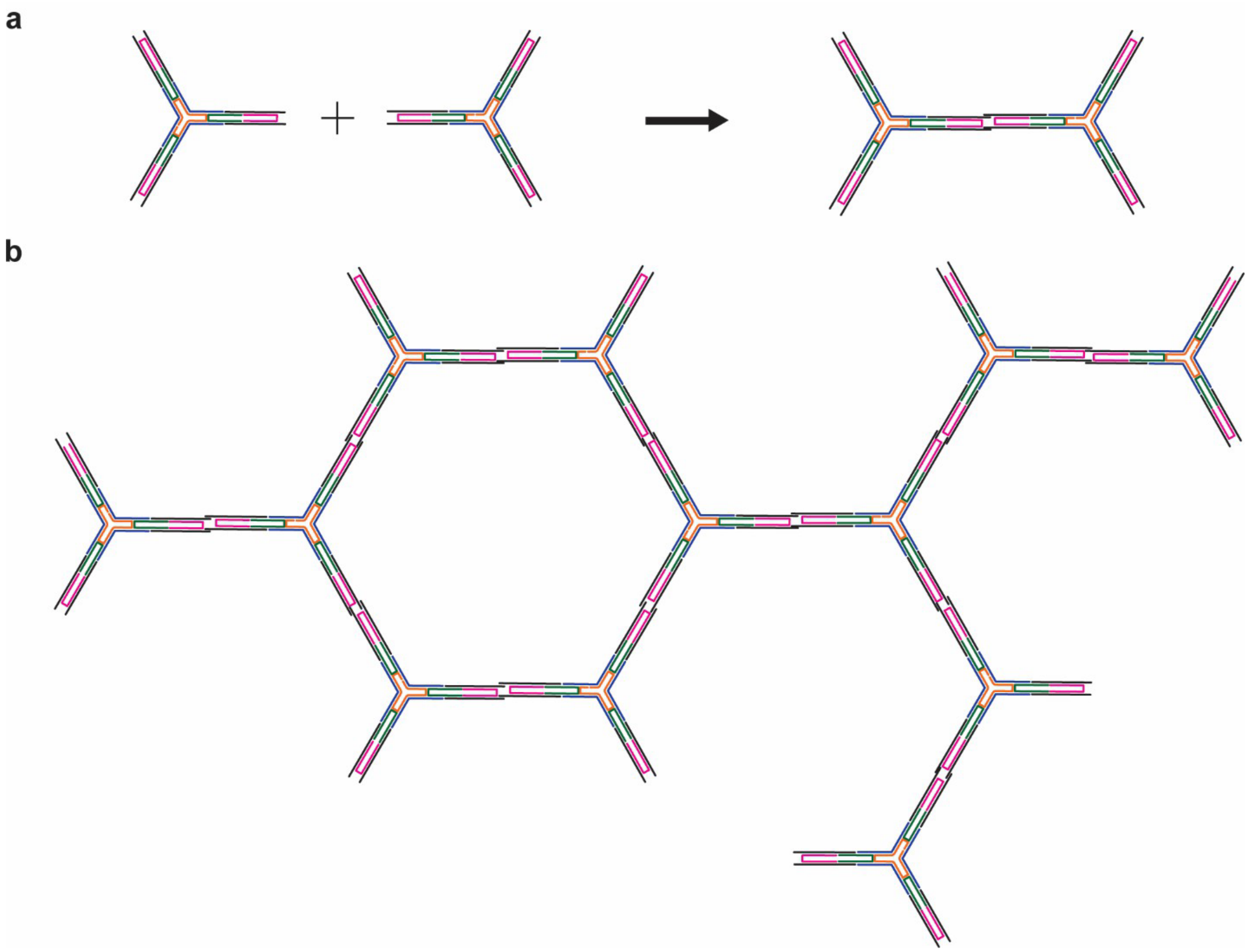


**Figure S2. Schematic of the putative DNA hydrogel mesh formation with 3WJ structural motifs. a)** 3WJ regular arms motifs with DNA linkers (linkers of A motif hybridize to linkers of B motif) to form a dimer. **b)** Potential graphic representation of a multi-motifs network.

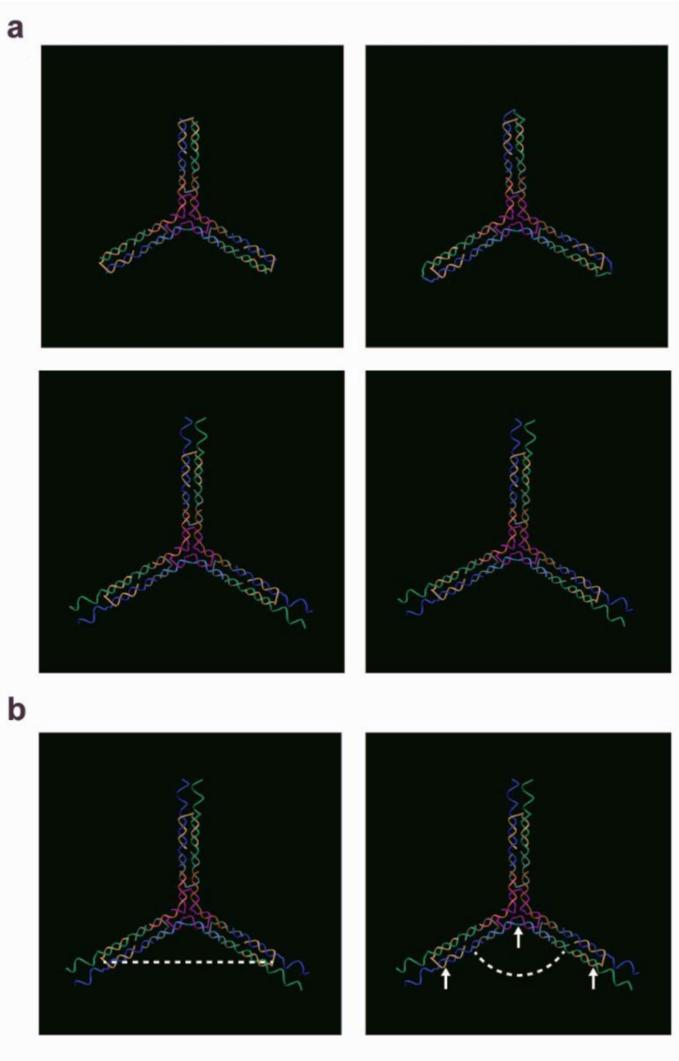


**Figure S3. Three dimensional representations and theoretical measurements of different 3WJ motif designs used in this study. a)** 3WJ motifs with Bl, Sh, Lo, and Lm linkers (*clockwise from top left*). **b)** Estimation of the distance (*left*) and angle (*right*) between each arm on the 3WJ motif, shown here with the Lo linker. White arrows denote specific atoms selected for measurements.

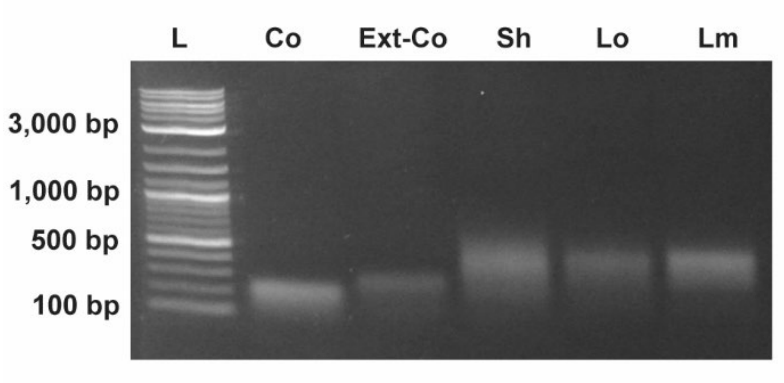


**Figure S4. Agarose gel characterization of the 3WJ motifs with regular arms from the core to the fully assembled structures with linkers.** Core (Co) and extended-core (Ext-Co) components shown along full motifs folded with short (Sh), long (Lo), and long mismatch (Lm) linker types.

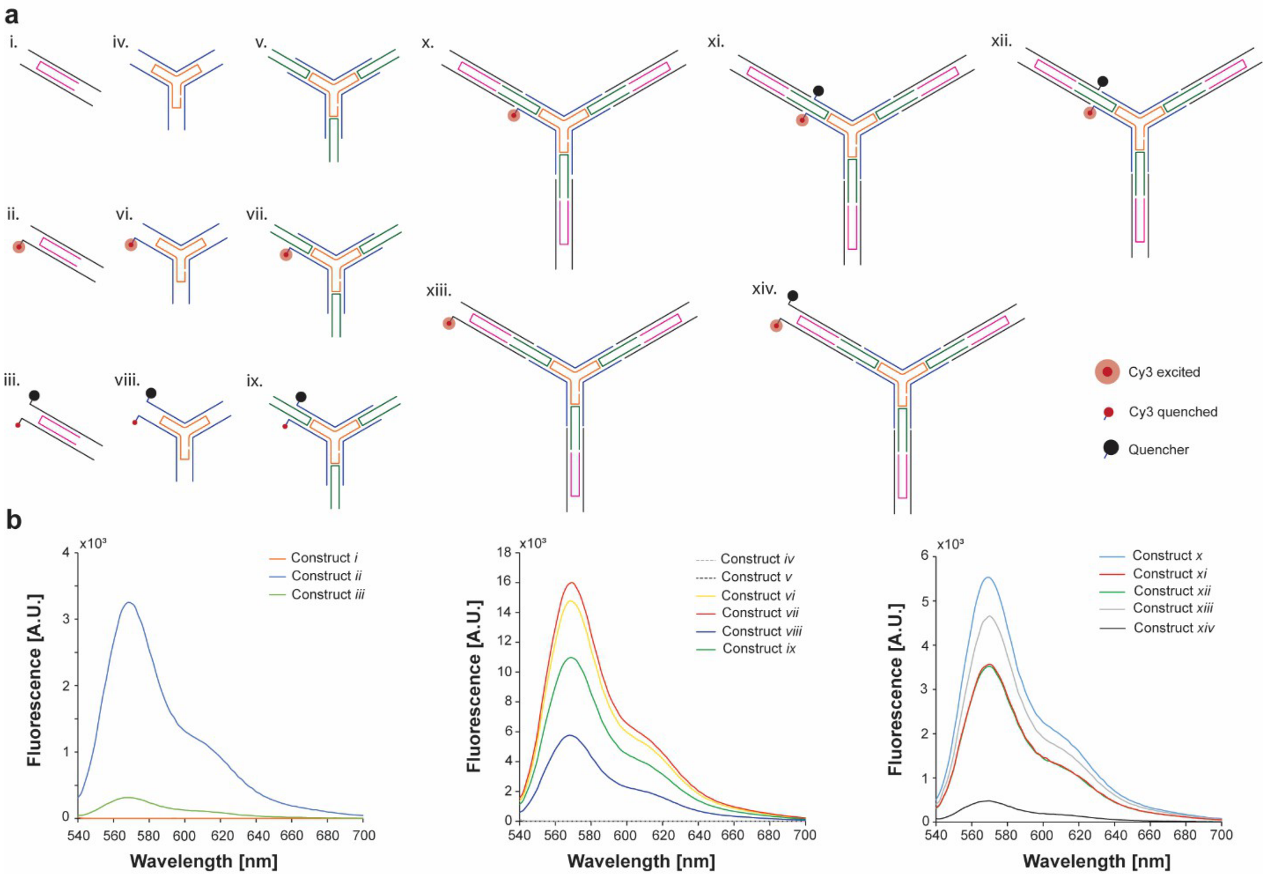


**Figure S5. FRET assay with Cy3/IowaBlack dye/quencher pair to monitor sequential formation of the DX-tile motifs. a)** schematic representation of the different constructs tested and the location of the FRET pairs. **b)** Fluorescence measurements of the different constructs after annealing. The constructs were excited at 520 nm wavelength and the emission was recorded from 540 nm to 700 nm.

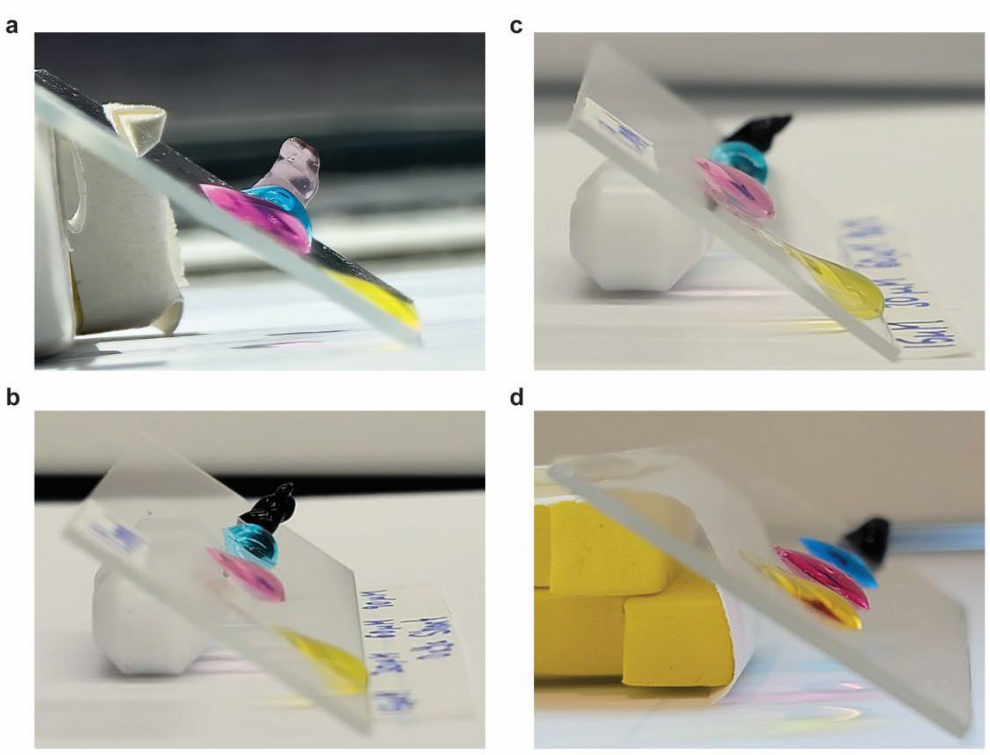

**Figure S6. Representative 3WJ gel array images on a tilted surface.** The 40 μL samples were made at 15 μM (yellow), 30 μM (pink), 60 μM (blue), and 90 μM (black). **a)** 3WJ-Bl. **b)** 3WJ-Sh. **c)** 3WJ-Lo. **d)** 3WJ-Lm.

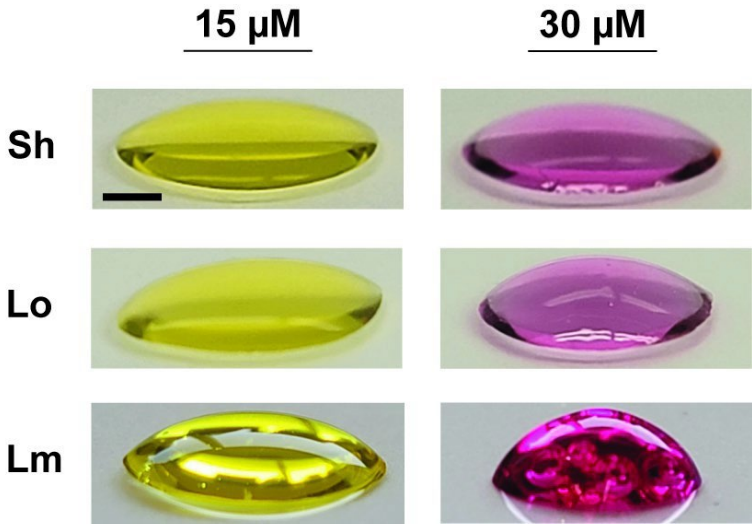


**Figure S7. Representative 3WJ gel array images at 15 and 30 μM.** All samples were made at 40 μL. (Scale bar: 5 mm).

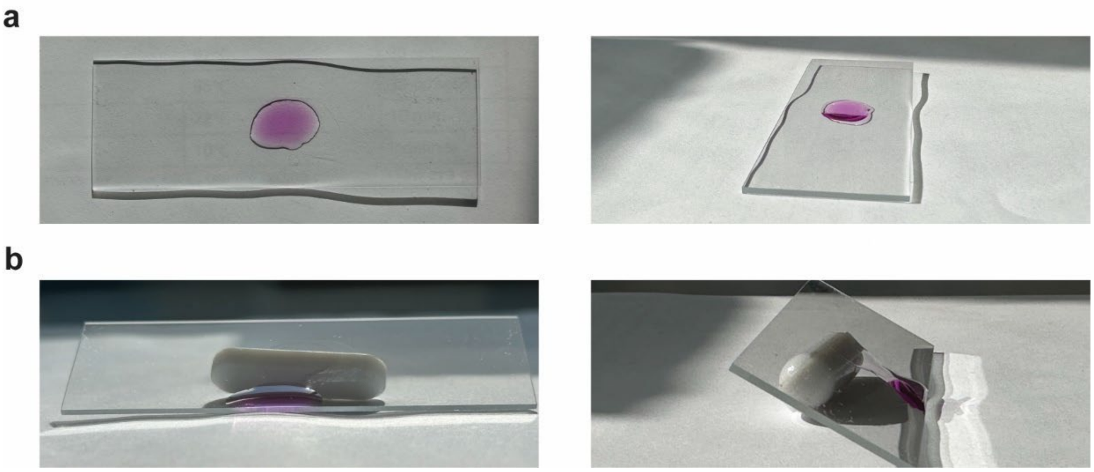


**Figure S8. 3WJ mixture with single motif type or noncomplementary motifs fail to form hydrogel.** Representative images of 3WJ-Lm-60μM motif B on **a)** flat and **b)** tilted surfaces. The two samples were made at 40 μL.

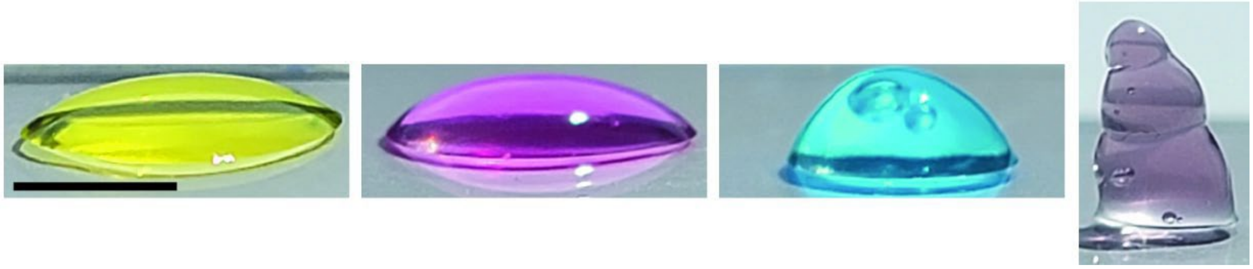

**Figure S9. Representative full gel array images of 3WJ-Bl hydrogel.** From left to right: 15 μM, 30 μM, 60 μM, and 90 μM (Scale bar: 5 mm). All sample were made at 40 μL.

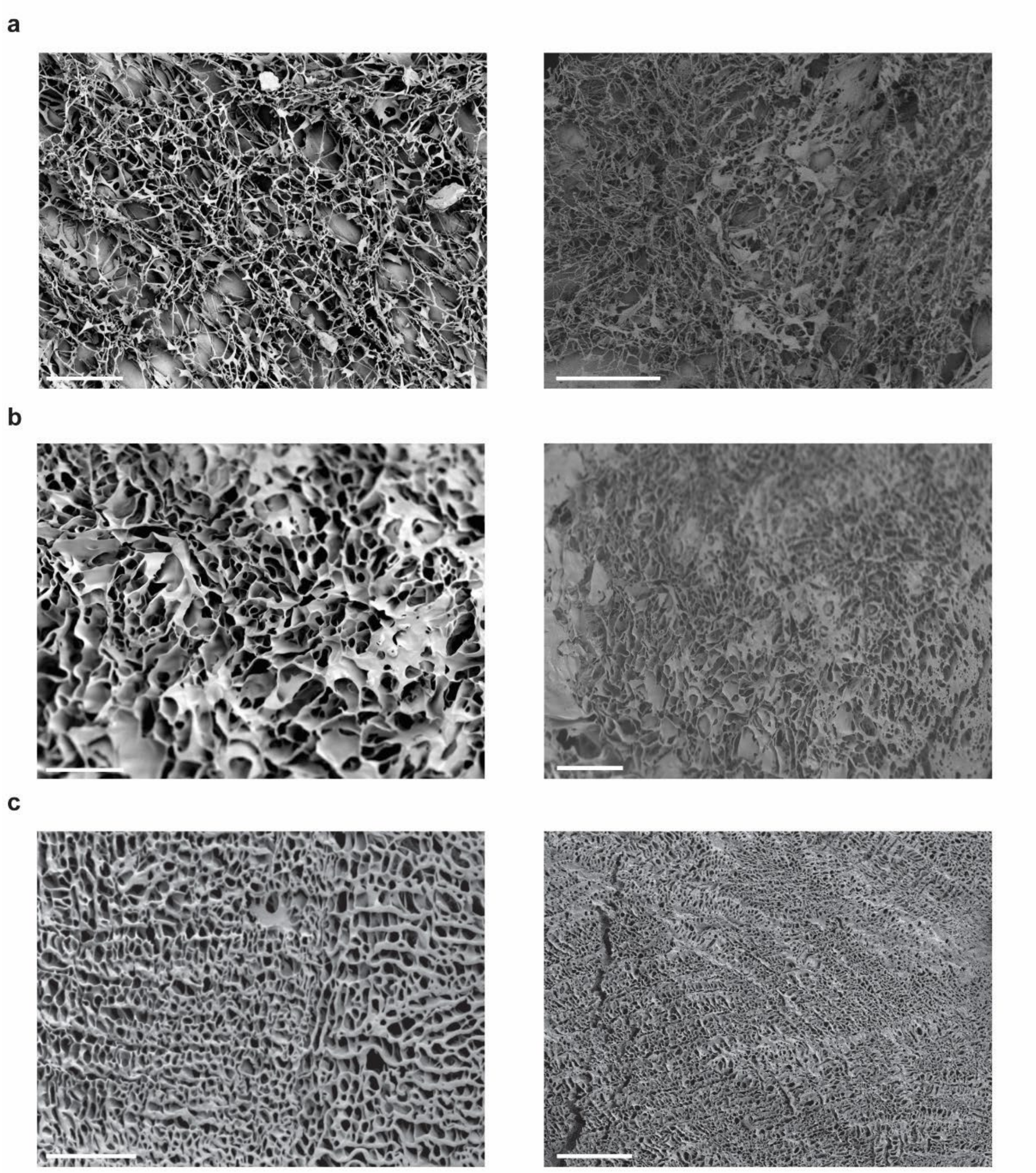


**Figure S10. Representative (*left*; scale bar: 50 μm) and full (*right*; scale bar: 100 μm) scanning electron microscopy (SEM) images of 3WJ-Bl hydrogel. a)** 30 μM, **b)** 60 μM, and **c)** 90 μM.

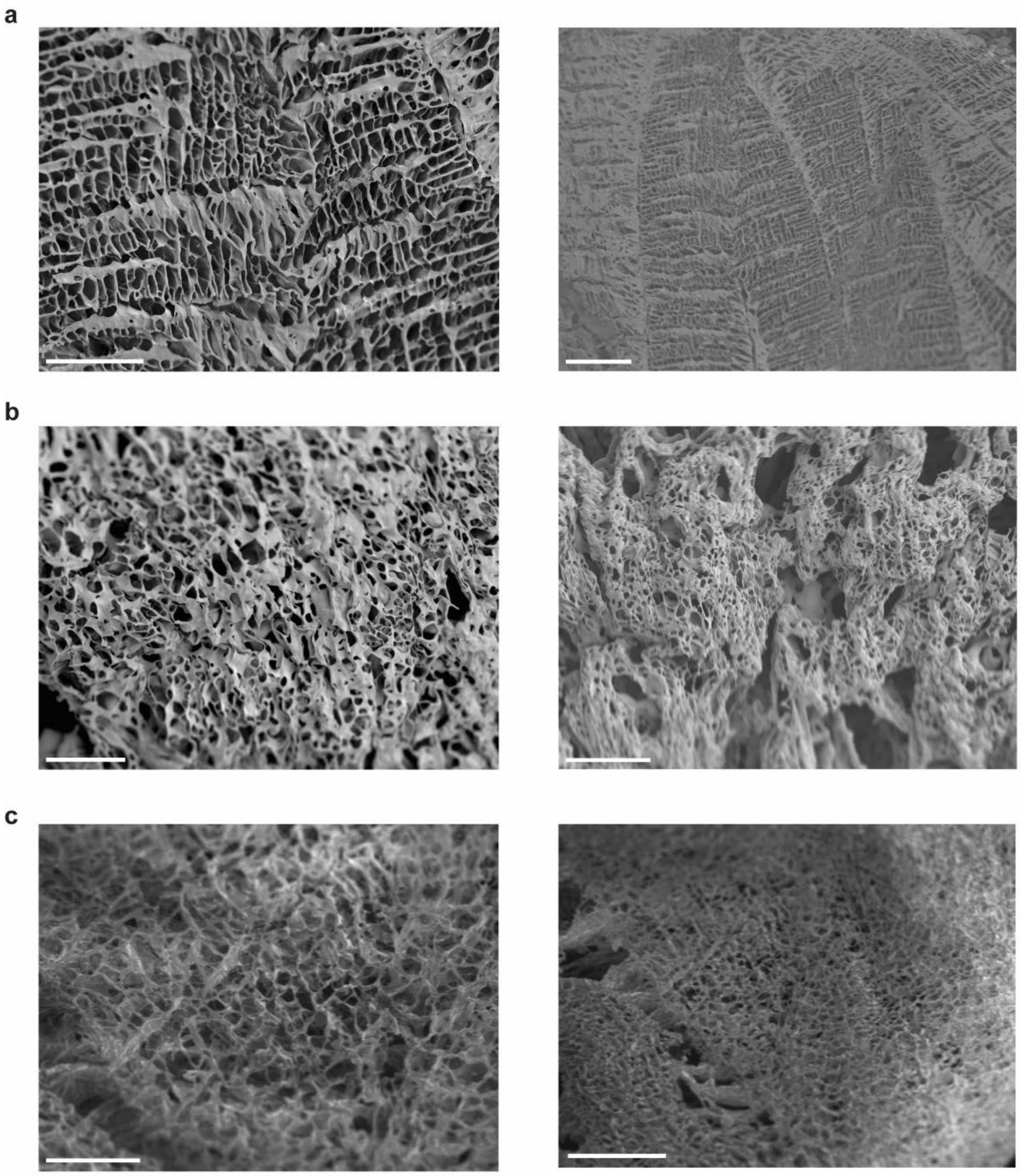


**Figure S11. Representative (*left*; scale bar: 50 µm) and full (*right*; scale bar: 100 µm) SEM images of 3WJ hydrogels at 30 µM. a)** Sh, **b)** Lo, and **c)** Lm linkers.

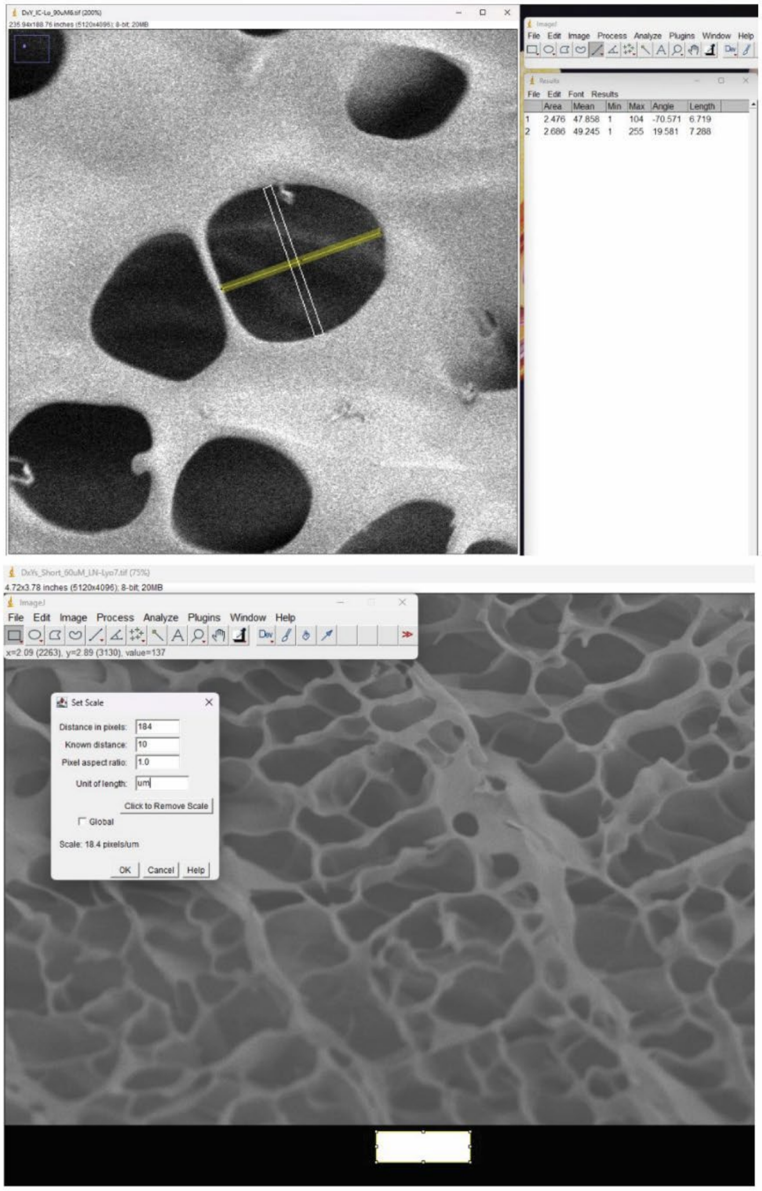


**Figure S12. Pore size measurement procedure used for SEM images of DX-tile hydrogels.**

Protocol followed for pore size measurements using Image J and described in Methods.

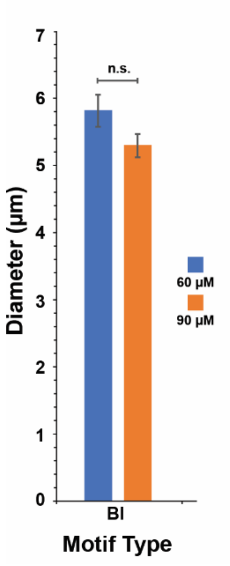


**Figure S13. Average pore size diameter for 3WJ-Bl hydrogels.** Error bars represent standard deviation of the mean (n = 50 independent samples/group) and p values were calculated from a student t-test (not significant, n.s.).

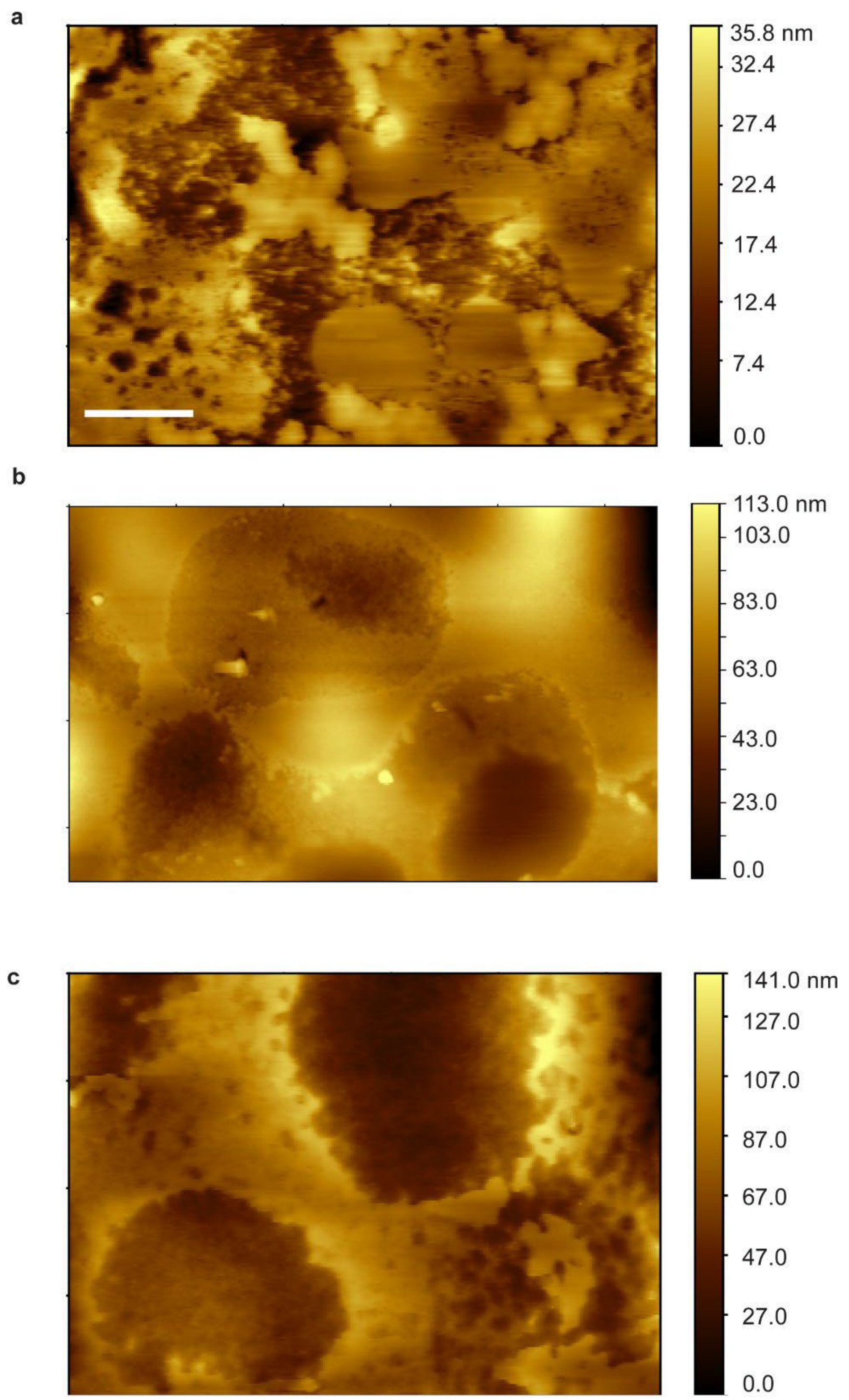


**Figure S14**. **Full-size AFM images of vacuum-dried 3WJ hydrogels. a)** Sh, **b)** Lo, and **c)** Lm (scale bar: 1 µm). All samples imaged at 60 µM.

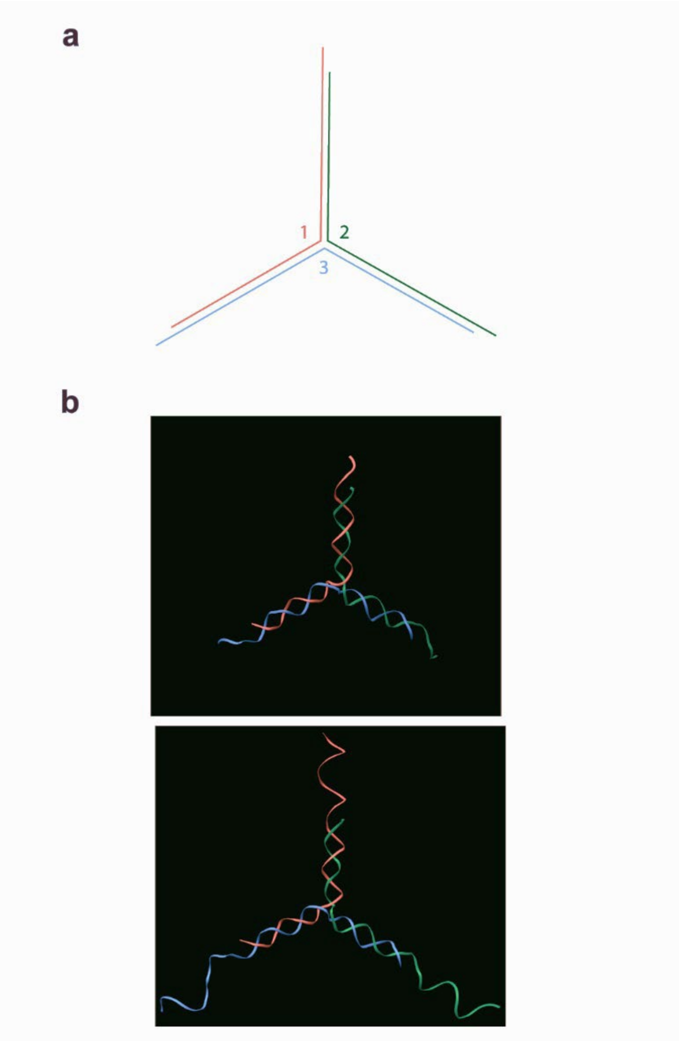


**Figure S15. Single duplex Y-motifs used in this study. a)** Numbering schematic. **b)** Three dimensional models of the Y7 (*top*) and Y21 (*bottom*) motifs.

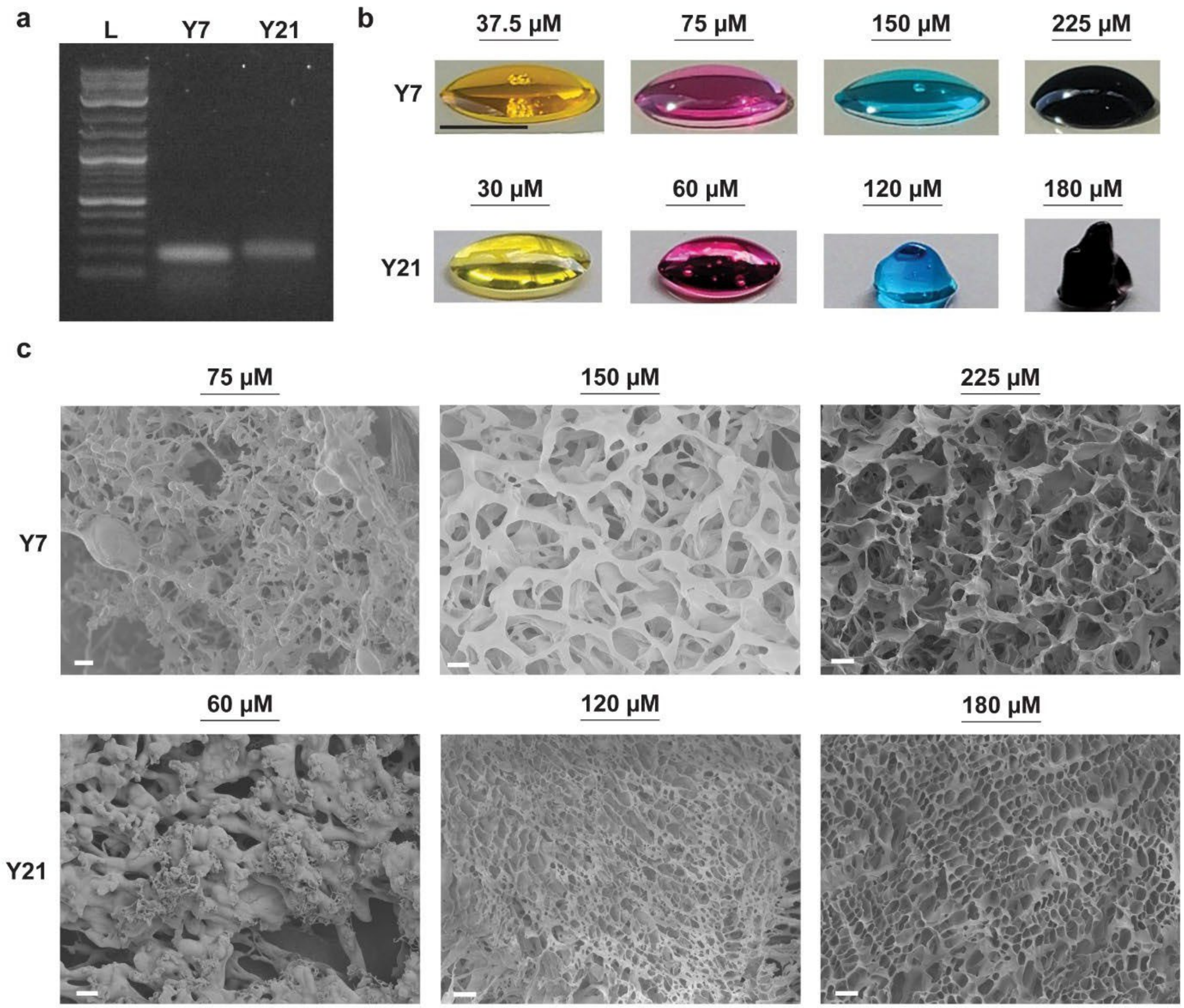


**Figure S16. Y motif folding and hydrogel formation. a)** Agarose gel electrophoresis confirming motif folding. **b)** Array images of hydrogels at comparable concentrations. **(**Scale bar: 5 mm). **c)** Representative SEM images verifying hydrogel formation. (Scale bars: 20 µm).

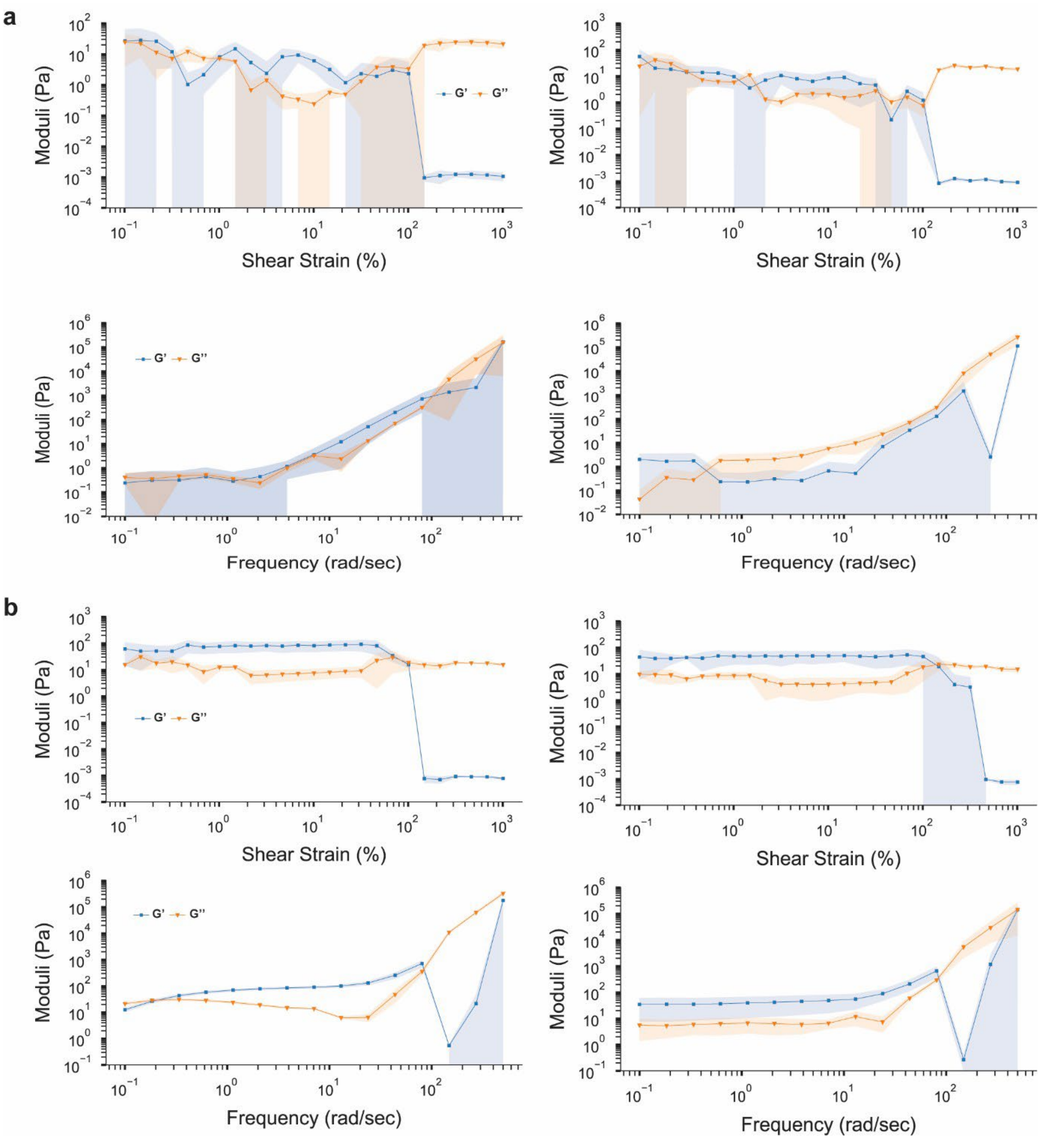


**Figure S17. Rheological profile of Y7 and Y21 hydrogels. a)** Amplitude (*top*) and frequency sweeps (*bottom*) of Y7 (*left*) and Y21 (*right*) hydrogels at 60 µM. Error shadings represent standard deviation of the mean (n = 3 independent samples/group). **b)** Amplitude (*top*) and frequency sweeps (*bottom*) of Y7 (*left*) and Y21 (*right*) hydrogels at 150 and 120 µM, respectively. Error shadings represent standard deviation of the mean (n = 3 independent samples/group).

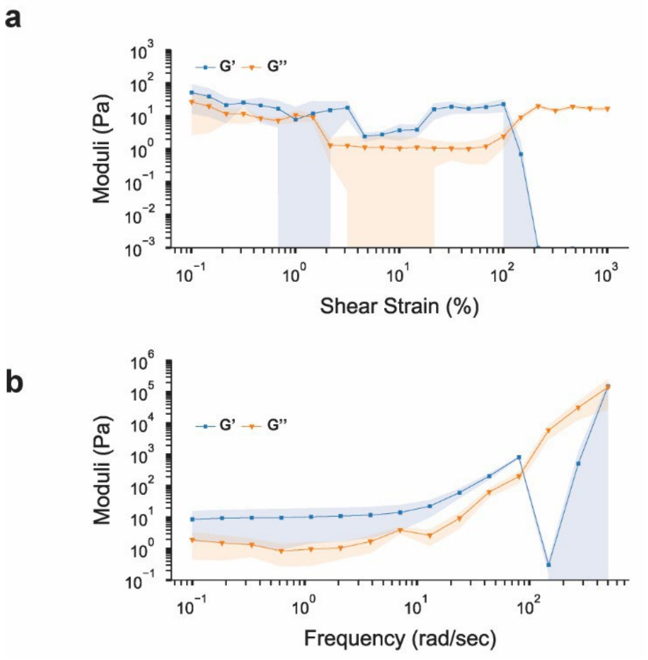


**Figure S18. Rheological profile of single DX-tile motif (3WJ-Sh, 60 µM)**. **a)** Amplitude sweeps. Error shadings represent standard deviation of the mean (n = 3 independent samples/group). **b)** Frequency sweeps. Error shadings represent standard deviation of the mean (n = 3 independent samples/group).

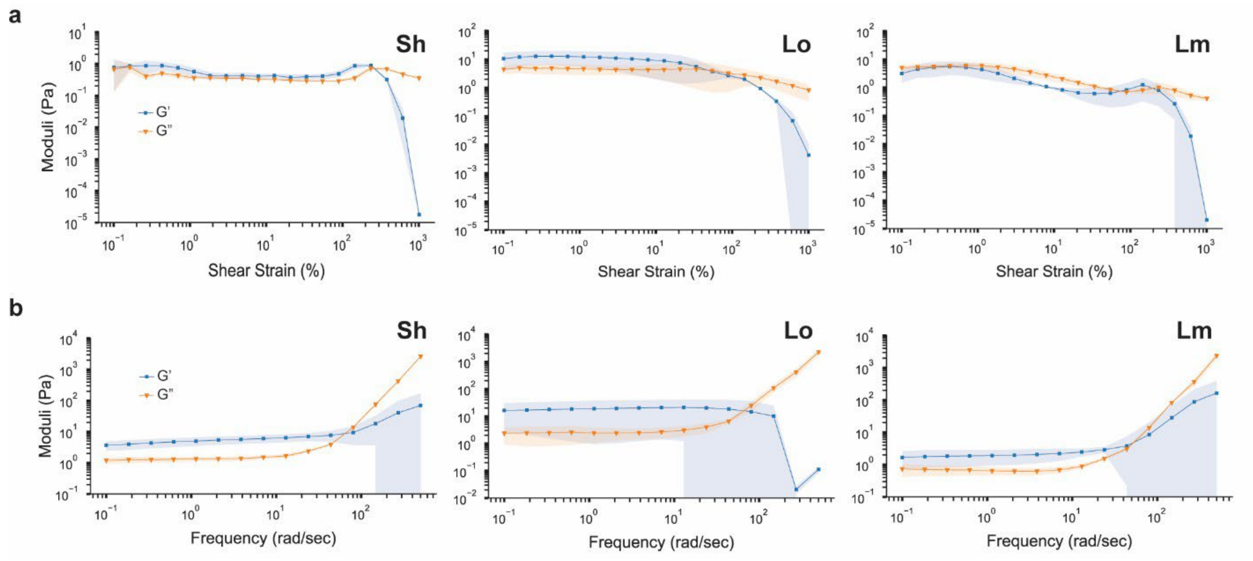


**Figure S19**. **Rheological characterization profile of the 3WJ constructs at 30 μM. a)** Amplitude sweeps. Error shadings represent standard deviation of the mean (n = 3 independent samples/group). **b)** Frequency sweeps. Error shadings represent standard deviation of the mean (n = 3 independent samples/group).

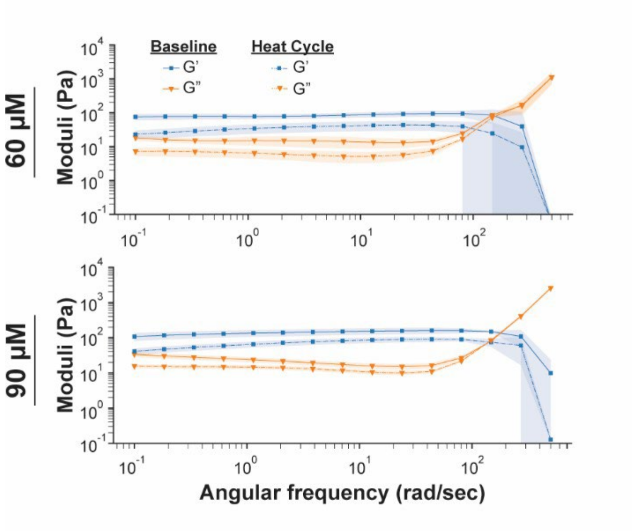


**Figure S20**. **Rheological characterization profile of the 3WJ hydrogel pre- and post- heat cycling**. Frequency sweeps of 3WJ-Sh hydrogel performed for 60 (top) µM and 90 µM (bottom). Error shadings represent standard deviation of the mean (n = 3 independent samples/group).

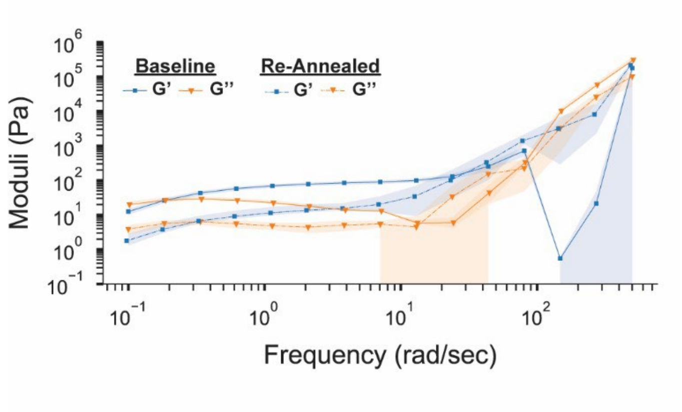


**Figure S21. Rheological characterization profile of the Y7 hydrogel pre- and post- heat cycling**. Frequency sweep of Y7 hydrogel (150 μM). Error shadings represent standard deviation of the mean (n = 3 independent samples/group).

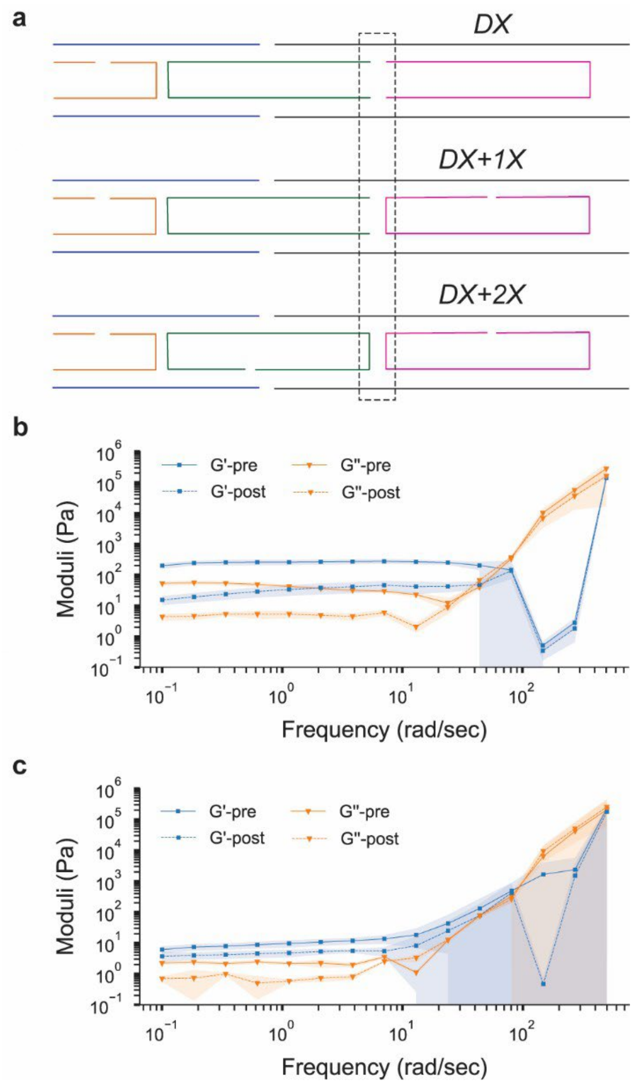


**Figure S22**. **Design schematic and rheological characterization profile of the 3WJ with the single and double crossovers**. **a)** Schematic of the original DX arm (*top*) compared to that with one additional crossover (*1X; middle*) and two additional crossovers (*2X; bottom*). **b)** Frequency sweeps of 3WJ-Sh-1X-60µM pre- and post-heat cycle. Error shadings represent standard deviation of the mean (n = 3 independent samples/group). **c)** Frequency sweeps of 3WJ-Sh-2X-60µM with two additional crossovers in the arms pre- and post-heat cycle. Error shadings represent standard deviation of the mean (n = 3 independent samples/group).

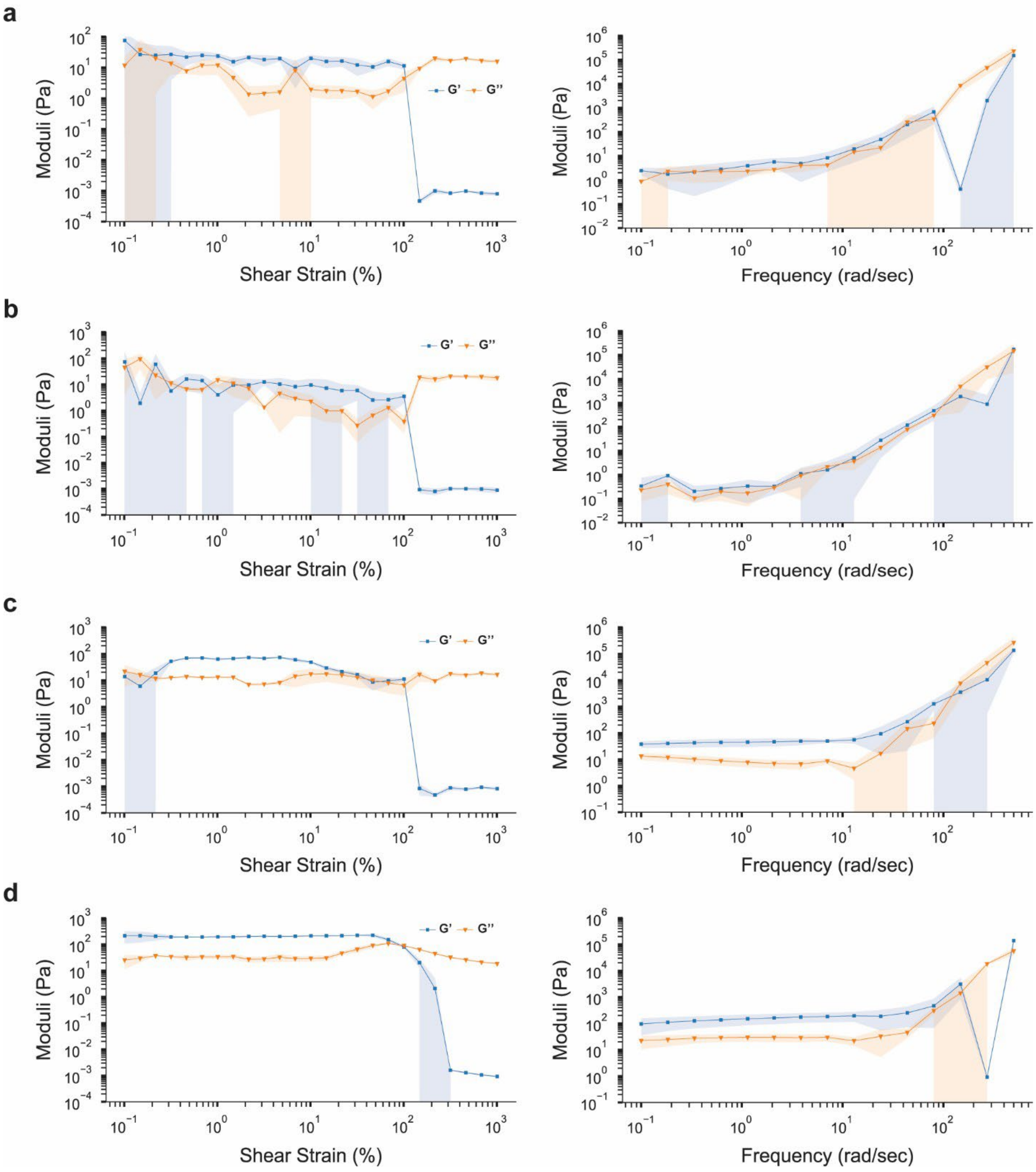


**Figure S23**. **Rheological characterization profile of the 3WJ with the short and long arms**. All samples were made with Sh linker at 60 μM. **a)** Short arm amplitude sweep (*left*) and frequency sweep (*right*). Error shadings represent standard deviation of the mean (n = 3 independent samples/group). **b)** Short arm-open nick amplitude sweep (*left*) and frequency sweep (*right*). Error shadings represent standard deviation of the mean (n = 3 independent samples/group). **c)** Long arm amplitude sweep (*left*) and frequency sweep (*right*). Error shadings represent standard deviation of the mean (n = 3 independent samples/group).

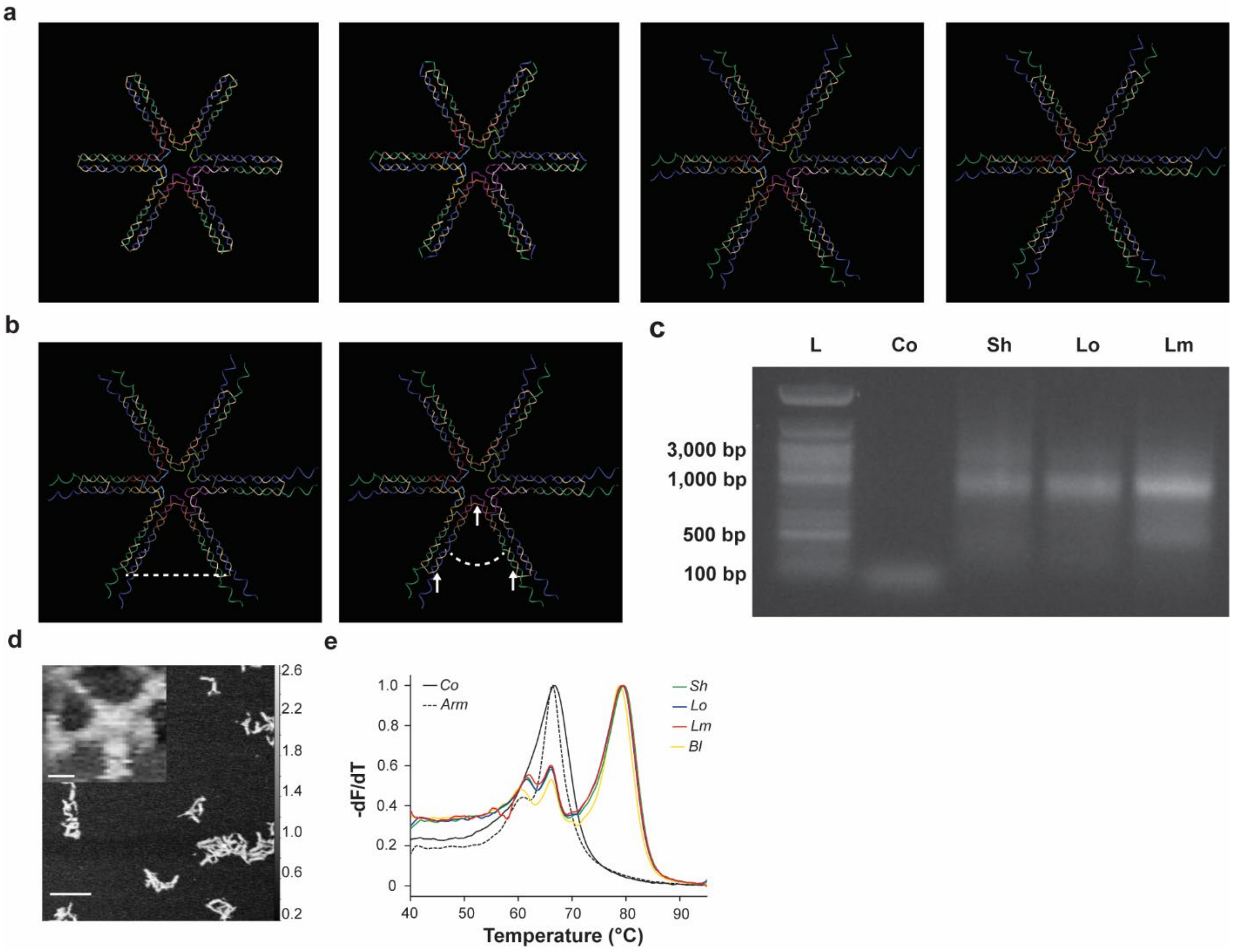


**Figure S24. Characterization profile of the 6WJ constructs**. **a)** 3D representation of 6WJ motifs with Bl, Sh, Lo, and Lm linkers (*left to right*). **b)** Estimation of the distance (*left*) and angle (*right*) between each arm on the 6WJ motif, shown here with the Lo linker. White arrows denote specific atoms selected for measurements. **c)** Agarose gel characterization of Core (Co) component shown along motifs folded with Sh, Lo, and Lm linker types. **d)** AFM showing folding of individual motif (full scale bar: 100 nm; inset scale bar: 10 nm). e**)** Melting curve of motif constituents (n = 4 independent samples/group).

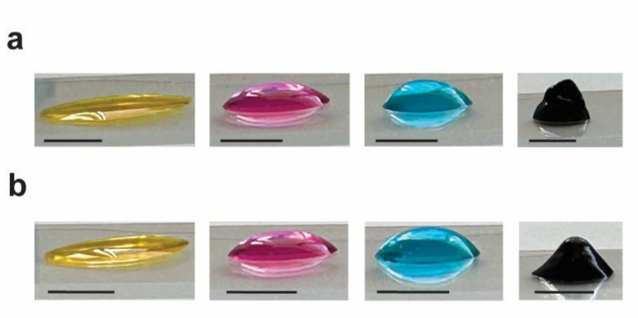


**Figure S25. Representative array images of 6WJ hydrogels.** From left to right: 7.5 μM, 15 μM, 30 μM, and 45 μM hydrogels formed with **a)** Sh and **b)** Lo linkers (Scale bar: 5 mm). All sample were made at 40 μL.

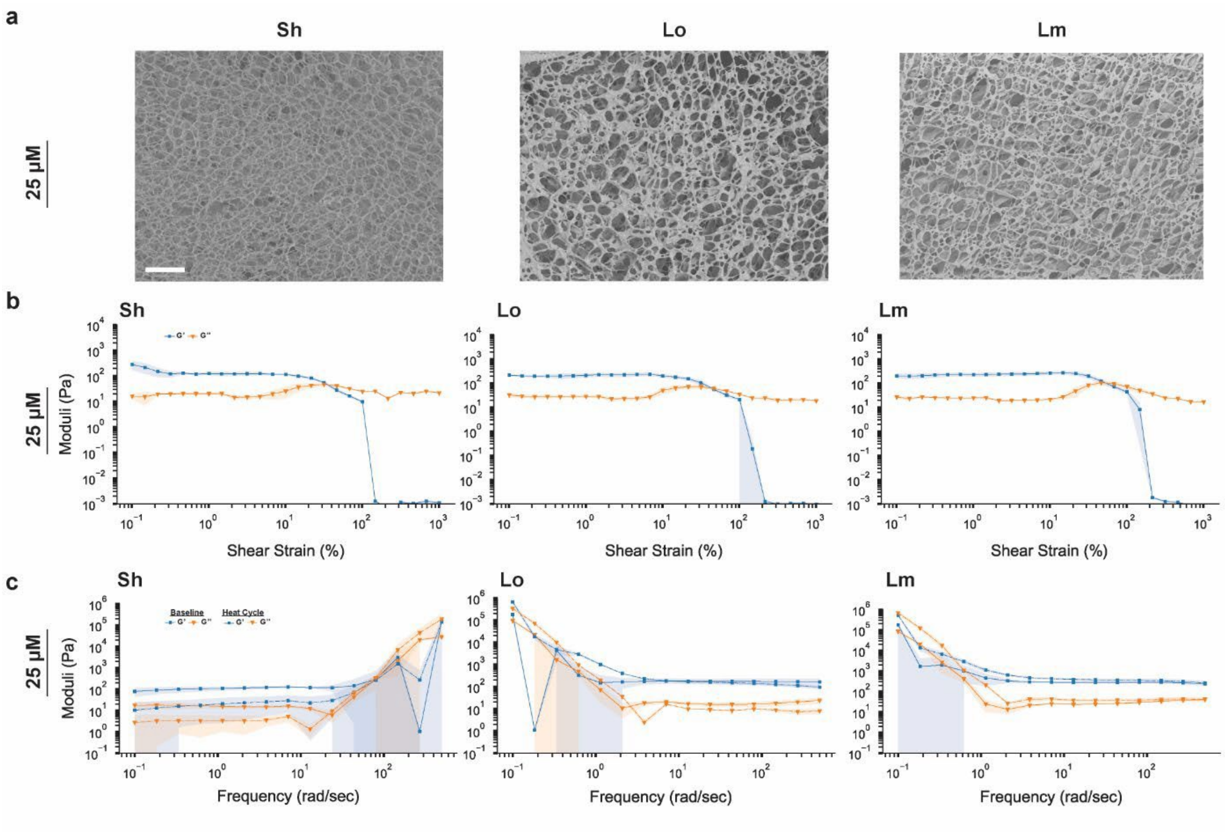


**Figure S26. Characterization profile of the 6WJ hydrogels at 25 µM with Sh, Lo, and Lm linkers** (*left to right*). **a)** Representative SEM images of 6WJ hydrogels (scale bar: 50 µm). **b)** Amplitude sweeps. Error shadings represent standard deviation of the mean (n = 3 independent samples/group). **c)** Frequency sweeps. Error shadings represent standard deviation of the mean (n = 3 independent samples/group).

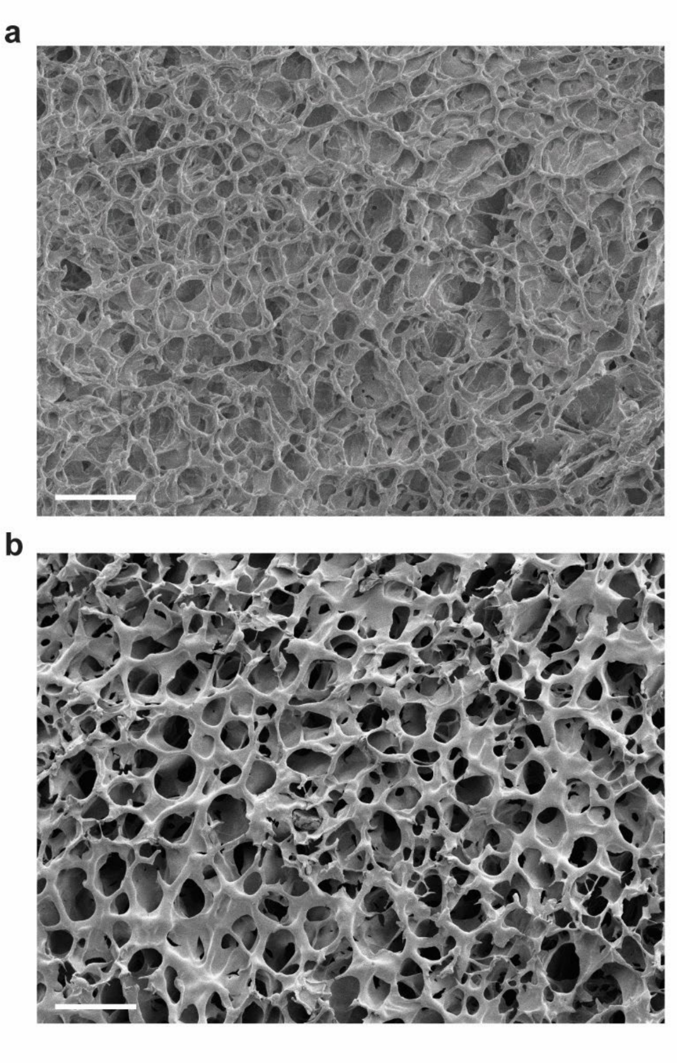


**Figure S27.** Representative scanning electron microscopy (SEM) images of 6WJ-Bl hydrogel at **a)** 25 µM and **b)** 50 µM. (scale bar: 50 µm)

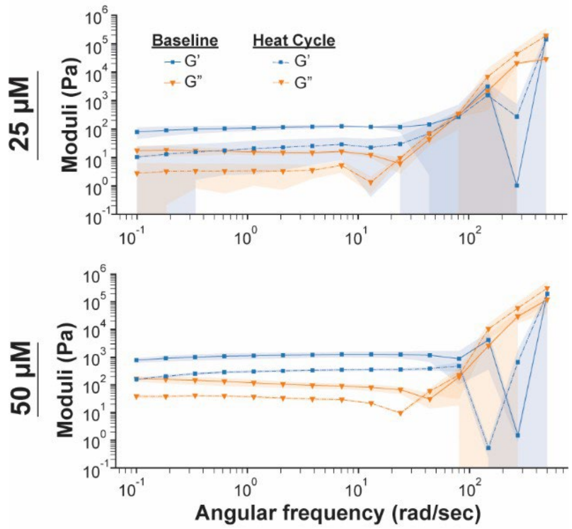


**Figure S28**. **Rheological characterization profile of the 6WJ hydrogel pre- and post- heat cycling**. Frequency sweeps of 6WJ-Sh hydrogel performed for 25 μM (top) and 50 μM (bottom). Error shadings represent standard deviation of the mean (n = 3 independent samples/group).

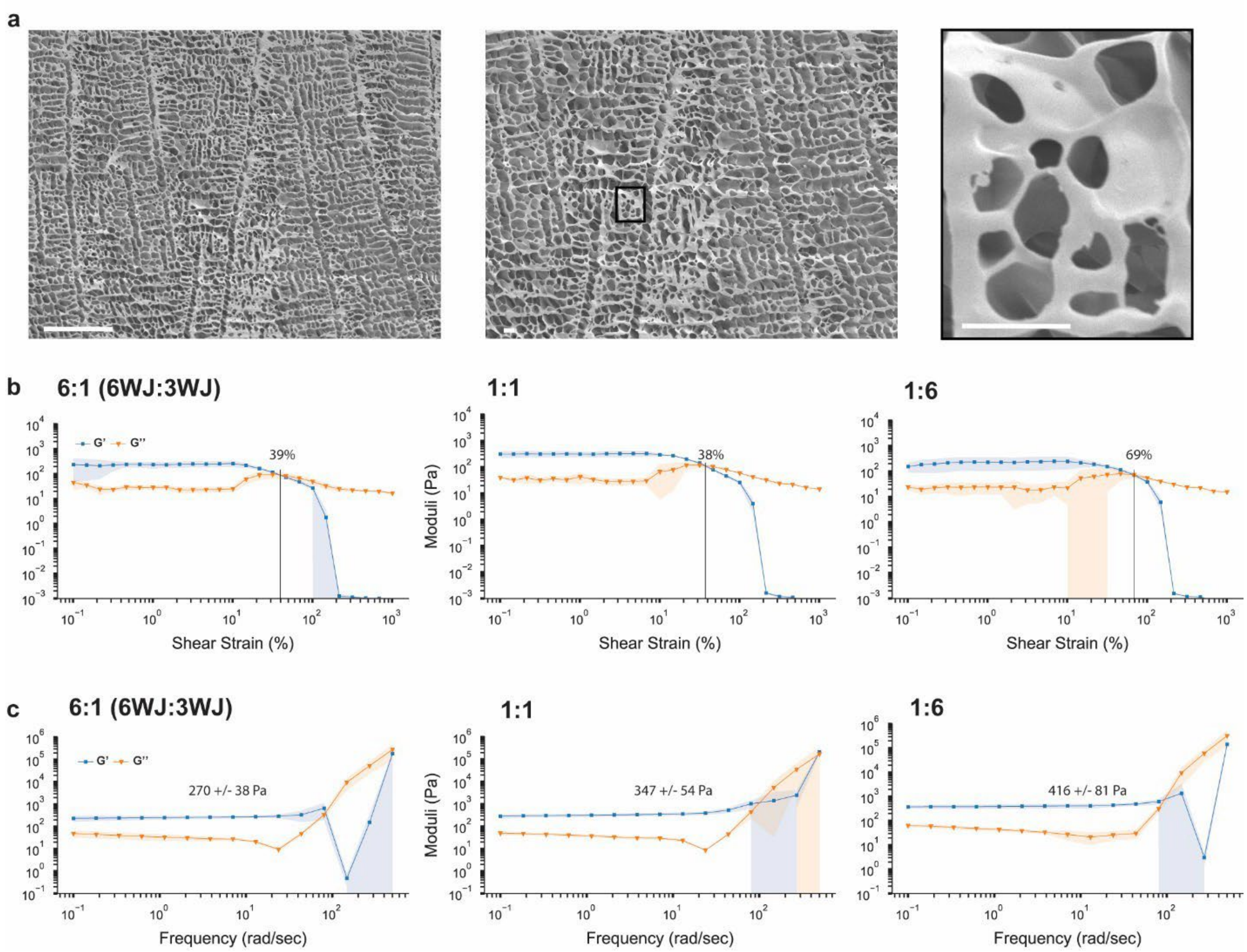


**Figure S29**. **SEM and Rheological characterization profile of the combination 6WJ:3WJ hydrogels**. **a)** Representative SEM images of 6WJ:3WJ hydrogels at 40:80 µM ratio. Scale bars: 100 µm (*left*) and 10 µm (*middle*, *right*). **b)** Amplitude sweeps of 6WJ:3WJ hydrogels at 30:60 µM ratio. Error shadings represent standard deviation of the mean (n = 3 independent samples/group). **c)** Frequency sweeps of 6WJ:3WJ hydrogels at 30:60 µM ratio. Error shadings represent standard deviation of the mean (n = 3 independent samples/group).

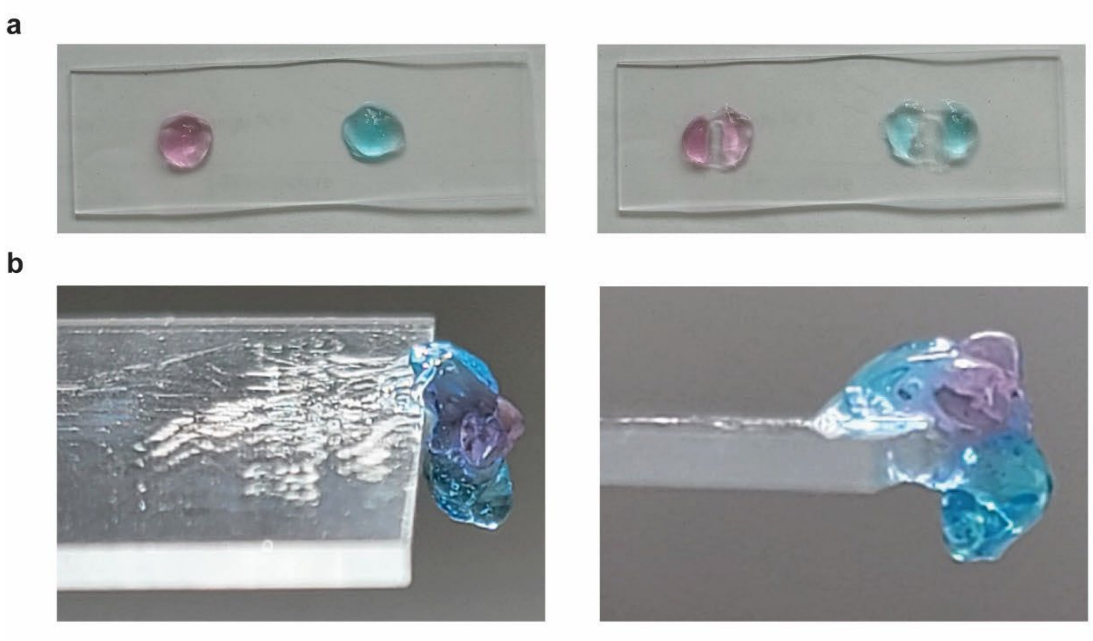

**Figure S30. Demonstration of hydrogel self-healing**. Representative images of 3WJ-Lo-90uM hydrogel above show: **a)** initial formation and coloring of two separate gels (left) and the initial cut (right). **b)** combination of hydrogel halves to form new hydrogel hanging off edge of standard microscope slide.

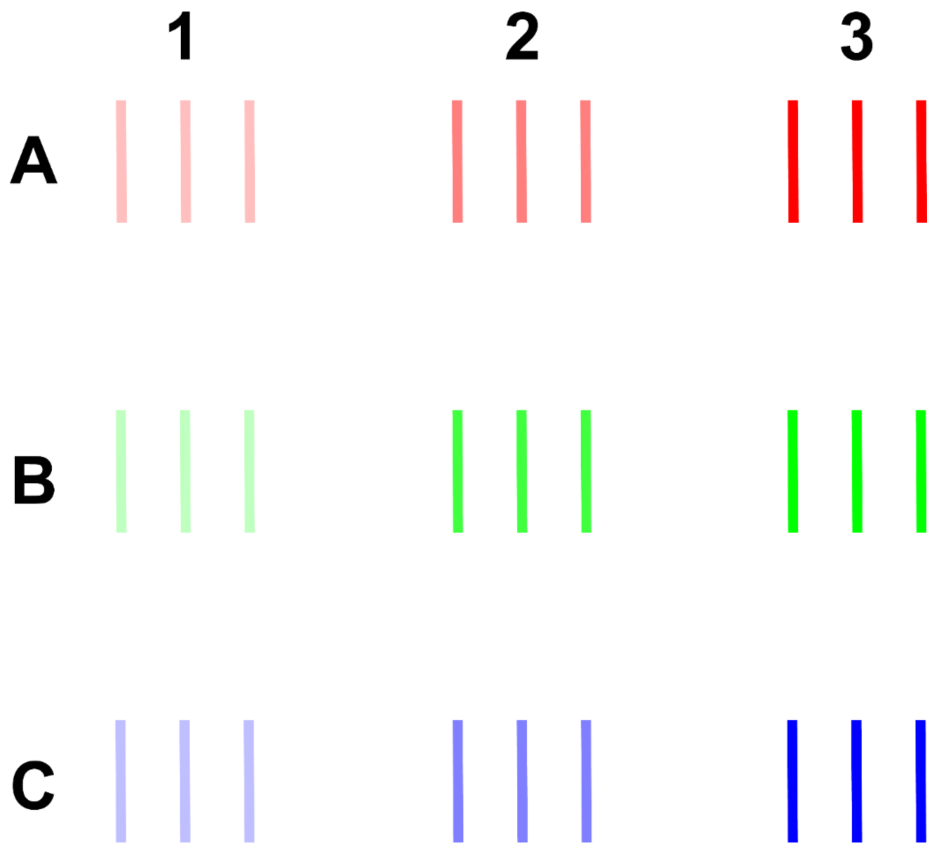


**Figure S31. Line test print.** Initial line tests performed with different parameters, such as line thickness (1X inner diameter of needle, A; 2X inner diameter of needle, B; 3X inner diameter of needle, C), print speed (6 mm/s, 1; 9 mm/s, 2; 12 mm/s, 3), needle gauge (25G and 30G; not shown), and applied pressure (30 PSI, 35 PSI, and 40 PSI).

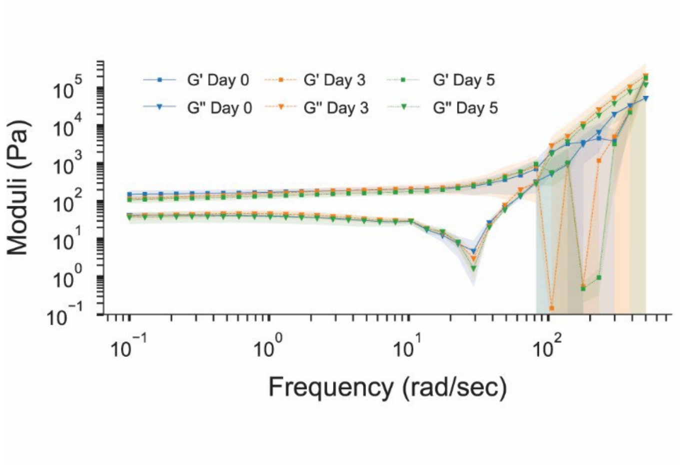


**Figure S32. Stability of elastic modulus for 3WJ-Sh hydrogels (60 µM) as evaluated by rheology**. Frequency sweep of gels over five days of storage (Day 1, Day 3, Day 5) following folding.
